# In silico karyotyping of chromosomally polymorphic malaria mosquitoes in the *Anopheles gambiae* complex

**DOI:** 10.1101/687566

**Authors:** R. Rebecca Love, Seth N. Redmond, Marco Pombi, Beniamino Caputo, Vincenzo Petrarca, Alessandra della Torre, The *Anopheles gambiae* 1000 Genomes Consortium, Nora J. Besansky

**Affiliations:** Eck Institute for Global Health & Department of Biological Sciences, University of Notre Dame, Notre Dame, IN 46556, USA; Infectious Disease and Microbiome Program, Broad Institute, Cambridge, MA 02142, USA; Dipartimento di Sanità Pubblica e Malattie Infettive, Istituto Pasteur Italia-Fondazione Cenci-Bolognetti, Università di Roma “La Sapienza”, Piazzale Aldo Moro, 5, 00185 Rome, Italy

**Author notes:** https://www.malariagen.net/projects/ag1000g#people (The *Anopheles gambiae* 1000 Genomes Consortium). Corresponding author, **Corresponding Author:** Nora J. Besansky, Department of Biological Sciences, University of Notre Dame, Notre Dame, IN 46556 USA. Monash University, Institute of Vector-Borne Disease, 3800, Clayton, Australia (Seth N. Redmond).

**Keywords:** *Anopheles gambiae*, chromosomal inversion polymorphism, genomics, inversion genotyping, karyotype analysis, malaria vector, tag SNP

## Abstract

Chromosomal inversion polymorphisms play an important role in adaptation to environmental heterogeneities. For mosquito species in the *Anopheles gambiae* complex that are significant vectors of human malaria, paracentric inversion polymorphisms are abundant and are associated with ecologically and epidemiologically important phenotypes. Improved understanding of these traits relies on determining mosquito karyotype, which currently depends upon laborious cytogenetic methods whose application is limited both by the requirement for specialized expertise and for properly preserved adult females at specific gonotrophic stages. To overcome this limitation, we developed sets of tag SNPs inside inversions whose biallelic genotype is strongly correlated with inversion genotype. We leveraged 1,347 fully sequenced *An. gambiae* and *Anopheles coluzzii* genomes in the Ag1000G database of natural variation. Beginning with principal components analysis (PCA) of population samples, applied to windows of the genome containing individual chromosomal rearrangements, we classified samples into three inversion genotypes, distinguishing homozygous inverted and homozygous uninverted groups by inclusion of the small subset of specimens in Ag1000G that are associated with cytogenetic metadata. We then assessed the correlation between candidate tag SNP genotypes and PCA-based inversion genotypes in our training sets, selecting those candidates with >80% agreement. Our initial tests both in held-back validation samples from Ag1000G and in data independent of Ag1000G suggest that when used for *in silico* inversion genotyping of sequenced mosquitoes, these tags perform better than traditional cytogenetics, even for specimens where only a small subset of the tag SNPs can be successfully ascertained.

## Introduction

A chromosomal inversion originates when a chromosome segment reverses end to end. Inversions maintained in plant and animal populations as structural polymorphisms tend to be large (several megabases) and contain hundreds of genes (reviewed in Wellenreuther and Bernatchez 2018). Long-term balancing selection can maintain these polymorphisms through millions of generations and multiple species radiations (Wellenreuther and Bernatchez 2018). Because recombination is greatly reduced between opposite orientations in inversion heterozygotes, inversions preserve selectively advantageous combinations of alleles despite homogenizing gene flow in collinear regions. Theory and mounting evidence implicate inversions in local adaptation, adaptive divergence, and range expansion, though the precise molecular mechanisms are rarely known (Hoffmann *et al*. 2004; Kirkpatrick and Barton 2006; Hoffmann and Rieseberg 2008; Schaeffer 2008; Kirkpatrick 2010; Lowry and Willis 2010; Joron *et al*. 2011; Jones *et al*. 2012; Kirkpatrick and Barrett 2015; Twyford and Friedman 2015; Kapun *et al*. 2016; Ayala *et al*. 2017; Fuller *et al*. 2017; Wellenreuther *et al*. 2017; Wellenreuther and Bernatchez 2018). Importantly, because of occasional double-crossovers and gene conversion, the suppression of gene flux is not absolute. As long as inversion heterozygotes are formed in populations, any significant association between an inversion and an allele within its boundaries is subject to eventual erosion unless gene flux is countered by selection (Navarro *et al*. 1997; Andolfatto *et al*. 2001).

The *Anopheles gambiae* complex is a medically important group of at least eight closely related and morphologically indistinguishable mosquito sibling species from sub-Saharan Africa (White *et al*. 2011; Coetzee *et al*. 2013). Three members of the complex (the eponymous *Anopheles gambiae, Anopheles coluzzii*, and *Anopheles arabiensis*) are among the most significant malaria vectors globally, responsible for a majority of the 435,000 malaria deaths in 2017 (World Health Organisation 2018). The ecological plasticity of these three species contributes greatly to their status as major human malaria vectors (Coluzzi *et al*. 2002). In contrast to the other five, these three species have wide distributions across diverse biomes of tropical Africa. Not coincidentally, they also segregate strikingly high numbers of paracentric inversion polymorphisms, which are implicated in adaptation to seasonal and spatial environmental heterogeneities related both to climatic variables and anthropogenic alterations of the landscape (Coluzzi *et al*. 1979; Bryan *et al*. 1982; Coluzzi *et al*. 1985; Toure *et al*. 1998; Manoukis *et al*. 2008; Costantini *et al*. 2009; Simard *et al*. 2009; Cheng *et al*. 2012; Ayala *et al*. 2014; Caputo *et al*. 2014; Ayala *et al*. 2017; Cheng *et al*. 2018). Some of these inversions also have been associated with ecologically relevant phenotypes, including desiccation and thermal tolerance (Gray *et al*. 2009; Rocca *et al*. 2009; Cassone *et al*. 2011; Fouet *et al*. 2012; Ayala *et al*. 2018; Cheng *et al*. 2018).

The sister taxa *An. gambiae* and *An. coluzzii*, the focus of the present investigation, are the most closely related species in the *An. gambiae* complex, sharing extensive nucleotide variation through both recent common ancestry and introgression (Fontaine *et al*. 2015; Hanemaaijer *et al*. 2018), while maintaining characteristic differences in ecology and behavior (Costantini *et al*. 2009; Diabate *et al*. 2009; Simard *et al*. 2009; Gimonneau *et al*. 2010; Gimonneau *et al*. 2012a; Gimonneau *et al*. 2012b; Dabire *et al*. 2013; Tene Fossog *et al*. 2015; Ayala *et al*. 2017). They also share four of six common chromosomal inversion polymorphisms on chromosomal arm 2R (*b, c, d, u*) and the only inversion polymorphism on chromosomal arm 2L (*a*) (Figure 1) (della Torre *et al*. 2005). These inversions range in size from ~4Mb to 22Mb, and together span thousands of genes and a sizeable fraction of chromosome 2: ~61% of 2R and ~38% of 2L polytene (euchromatic) content (Coluzzi *et al*. 2002). Inversions 2L*a* and 2R*b* are found in populations throughout tropical Africa and are therefore cosmopolitan, while three other inversions on 2R (*c, d*, and *u*) are widespread in West, very rare in Central Africa, and absent from East Africa. The remaining two inversions, 2R*j* and 2R*k*, have more restricted geographic distributions (Coluzzi *et al*. 2002; Ayala *et al*. 2017).

**Figure 1.**
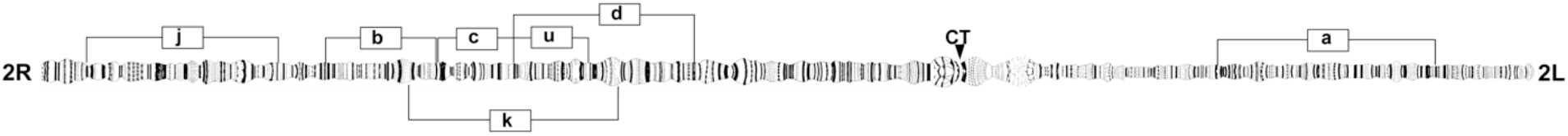
Diagrammatic representation of the common polymorphic inversions (labeled brackets) on chromosome 2 in *An. gambiae*. Polytene chromosome map modified from Figure 1 and poster in Coluzzi *et al*. (2002). CT, centromere.

Early cytogenetic studies of *An. gambiae* and *An. coluzzii*, presumed at the time to be a single heterogenous species, uncovered genetic discontinuities that led to the designation of five presumed assortatively-mating ‘chromosomal forms’: FOREST, SAVANNA, MOPTI, BAMAKO, and BISSAU (Coluzzi *et al*. 1985; Toure *et al*. 1998; Coluzzi *et al*. 2002; della Torre *et al*. 2005). They were delineated based on stable non-random associations of different sets of chromosome 2R inversions in co-occurring populations, and differed in larval ecology. Subsequent DNA-based studies identified fixed differences in the ribosomal DNA (rDNA), located in the pericentromeric region of the X chromosome, leading to the definition of two assortatively mating M and S ‘molecular forms’ of *An. gambiae* (della Torre *et al*. 2001). The molecular forms, which were eventually given specific status as *An. coluzzii* (formerly M) and *An. gambiae sensu stricto* (formerly S) (Coetzee *et al*. 2013), are incongruent with the chromosomal forms. Nearly all inversion associations segregate in both species albeit at different frequencies, and likely play similar roles in ecological specialization and adaptation in both *An. gambiae s.s*. (hereafter, *An. gambiae*) and *An. coluzzii* (della Torre *et al*. 2005; Costantini *et al*. 2009; Simard *et al*. 2009; Ayala *et al*. 2017). Hence, inversion associations are indicative of environmental heterogeneities more so than intrinsic reproductive boundaries.

Beyond a role in ecological specialization, inversions in the *An. gambiae* complex are also associated with vector traits affecting malaria transmission intensity and control: biting and resting behavior (Coluzzi *et al*. 1979; Riehle *et al*. 2017), seasonality (Rishikesh *et al*. 1985), morphometric variation (Petrarca *et al*. 1990), and *Plasmodium* infection rates (Petrarca and Beier 1992; Riehle *et al*. 2017). Although a robust molecular assay is available for genotyping inversion 2L*a* in natural populations (White *et al*. 2007), 2R inversions with characterized breakpoint sequences (*j, b, c*, and *u*) (Coulibaly *et al*. 2007a; Sangare 2007; Lobo *et al*. 2010) proved difficult to genotype molecularly at the breakpoints (Coulibaly *et al*. 2007b; Lobo *et al*. 2010), owing to extensive tracts of flanking repetitive DNA. The 2R*k* breakpoints have yet to be characterized, but recent localization of the 2R*d* breakpoints in the reference genome assembly using proximity-ligation sequencing (Corbett-Detig *et al*. 2019) also revealed high repeat content, suggesting that repetitive DNA at inversion breakpoints will pose a significant challenge both for breakpoint characterization and for molecular genotyping assays targeting breakpoint regions in these species.

Failure to account for the presence of inversions is a barrier to a more comprehensive understanding of epidemiologically relevant mosquito behavior and physiology. Inversion-blind analysis of population data can mislead population genetic inference, and create spurious associations in genome-wide association studies (Seich Al Basatena *et al*. 2013; Houle and Marquez 2015). Powerful genomic resources exist for *An. gambiae*, including a high-quality reference genome assembly (Holt *et al*. 2002) and a database of genomic variation (Ag1000G) based on deep genome re-sequencing of thousands of mosquitoes from natural populations across Africa (Miles *et al*. 2017). Unfortunately, inversion genotypes are not automatically revealed by genome re-sequencing, as reads are mapped to their position in the reference genome assembly, not their position in the re-sequenced mosquito genome. Despite advancing genome technology, the only method currently available to determine the *An. gambiae* karyotype is a method perfected half a century ago (Coluzzi 1968) involving cytological analysis of ovarian nurse cell polytene chromosomes (Coluzzi *et al*. 2002; Pombi *et al*. 2008). At best, such cytological analysis is severely rate-limiting because it is laborious and requires highly specialized training. At worst, it is prohibitive because it requires proper preservation of chromosomes harvested only from ovaries of adult females at a specific gonotrophic stage; suitable polytenization is absent at other gonotrophic stages as well as in males (della Torre 1997). While salivary glands of late fourth instar larvae also contain chromosomes with an adequate degree of polytenization, and the banding patterns of salivary and ovarian chromosomes are homologous in principle, most bands are difficult to homologize due to a different pattern of chromosome ‘puffing’ (della Torre 1997), rendering this alternative impractical. To overcome these impediments, our goal is to develop broadly accessible computational and molecular methods of genotyping chromosomal inversions in individual specimens of *An. gambiae* and *An. coluzzii*.

Here, we exploit the Ag1000G database and leverage the subset of cytologically karyotyped specimens within that database to develop a computational approach for karyotyping applicable to whole genome sequence data. We identify multiple tag single nucleotide polymorphisms (SNPs) significantly associated with inversions across geography that collectively predict with high confidence the genotypes of six common polymorphic inversions on chromosome 2 (*a, j, b, c, d, u*) in individually sequenced genomes of *An. coluzzii* and *An. gambiae*. We then apply this approach to data generated independently of Ag1000G to show that our approach has wider utility, even for specimens where only a small subset of the tag SNPs can be successfully ascertained.

## Methods

### Mosquito genotype data

Variant call data used for the discovery of inversion tag SNPs were accessed from Ag1000G (Miles *et al*. 2017) and Vector Observatory (VOBS; Table S1), projects of the Malaria Genomic Epidemiology Network (MalariaGEN; https://www.malariagen.net/) that provide catalogs of genomic sequence variation based on individual wild-collected *An. gambiae* and *An. coluzzii* mosquitoes sampled from multiple African countries and the Mayotte archipelago. With the exception of four atypical samples (see next section), we verified species identifications as reported in Ag1000G and VOBS using principal component analysis (PCA) of biallelic SNPs on the X chromosome. We excluded any specimens with more than 50,000 missing genotypes on chromosomal arm 2R (N=9), and any specimens subjected to whole genome amplification (WGA) prior to genomic sequencing (N=44), as PCA revealed strong biases associated with WGA. After filtering, we retained variant call data from 1,347 mosquitoes (Table S2).

### Karyotype imputation by local PCA

Cytological karyotype information derived from phase contrast microscopy of ovarian polytene chromosomes (della Torre 1997) was available only for a relatively small subset of specimens (N=373) in Ag1000G/VOBS (hereafter, Ag1000G for brevity). Thus, as a first step toward discovering SNPs putatively predictive of inversion status (tag SNPs), we imputed karyotypes computationally at each of six focal inversions (Figure 1), using local PCA (where ‘local’ refers to windows of the genome corresponding to chromosomal rearrangements). Ma and Amos (2012) showed that applying PCA to SNP genotypes in a window of the genome containing an inversion polymorphic in population genomic data (an approach that we call ‘PCA karyotyping’) produces a pattern of three equidistant clusters (stripes) in a plot of the first two principal components, assuming adequate numbers of each of three possible inversion genotypes: inverted and uninverted (standard) homokaryotypes, and heterokaryotypes. The two flanking stripes represent alternative homokaryotypes, and the middle stripe represents the inversion heterokaryotype, a 1:1 “admixture” between the two homokaryotype classes (Ma and Amos 2012).

To apply this approach, we combined specimens from both species (*An. gambiae* and *An. coluzzii*) and different geographic localities into a single metapopulation sample of 1,347 mosquitoes (Tables S1, S2). We identified a set of biallelic SNPs within inversion boundaries (Table S3) with potentially informative levels of polymorphism [minor allele count ≥3 and minimum alternate allele frequency (MAF) ≥0.15 for all inversions except 2R*d*, for which the MAF threshold was reduced to 0.03]. As 2R*d* overlaps 2R*u* in the genome (Figure 1), we limited consideration to only those SNPs found outside (proximal to) 2R*u* for PCA karyotyping of 2R*d* (Table S3). Next, we converted mosquito genotypes at these SNPs to a count of the number of alternate alleles (‘0’ if both matched the reference allele, ‘1’ or ‘2’ if one or both matched the alternate allele, respectively). Using the scikit-allel Python package v1.1.9 (Miles and Harding 2017), we then applied PCA to the resulting matrix of alternate allele counts, and represented the output as a scatter plot of the first two principal components for each mosquito in the population sample. The correct genotype corresponding to the two homokaryotype stripes was determined based on the inclusion in a given stripe of mosquitoes with cytologically determined karyotype. Based on this classification, mosquitoes without cytologically determined karyotypes were assigned a PCA karyotype.

The distinction between stripes was not always sharp; the stripes could be diffuse and oblique rather than tightly clustered. In extreme cases, stripes were not initially discernable. Through an iterative process of ‘leave one population sample out’ followed by PCA, we determined that absence of a clear three-stripe pattern was attributable to some or all of the same four atypical source populations, in particular, those from Kenya, Mayotte, The Gambia, and Guinea Bissau. The Kenyan sample has been found to display signs of extreme inbreeding (Miles *et al*. 2017), and Mayotte is an island whose mosquito population is plausibly subject both to inbreeding and a degree of isolation from mainland samples. The Gambia and Guinea Bissau are localities with unusually high degrees of hybridization and introgression between *An. gambiae* and *An. coluzzii* (Caputo *et al*. 2008; Oliveira *et al*. 2008; Caputo *et al*. 2011; Marsden *et al*. 2011; Weetman *et al*. 2012; Nwakanma *et al*. 2013). Where necessary, we removed these population samples, as well as two *An. gambiae*-*An. coluzzii* hybrid specimens from Burkina Faso and Guinea Conakry, and repeated the PCA. In addition, successful PCA karyotyping of 2R*d* and 2R*j* required the removal of all *An. coluzzii* specimens owing to taxonomic structuring of variation. Accordingly, PCA karyotyping was successful on all (2L*a*) or subsets (all 2R inversions) of the 1,347 specimens (Table S4).

### Discovery of SNPs predictive of inversion orientation

The PEST reference genome assembly for *An. gambiae* (AgamP4; Giraldo-Calderon *et al*. 2015) was derived from a colony whose karyotype was homozygous standard with respect to all common chromosomal inversions in this species. We therefore had the general expectation that an individual SNP might be a good predictor of chromosomal inversion orientation if the reference allele is strongly associated with the standard arrangement and the alternate allele is strongly associated with the inverted arrangement within and across population samples. As shown in Figure 2 in overview, we assessed SNP genotype-inversion genotype concordance for each inversion in individual mosquitoes, limiting our assessment to potentially more informative, higher frequency biallelic SNPs inside inversion boundaries (*i.e*., those with MAF≥5%). We converted both the SNP genotype and the corresponding mosquito’s PCA-based inversion genotype to single numbers, representing the count of alternate alleles (0, 1, or 2) in the case of SNP genotype, and the count of inverted chromosomes (0, 1, or 2) in the case of inversion genotype. Successfully performing tags are expected to have a SNP genotype that correlates strongly with the PCA-based inversion genotype.

**Figure 2.**
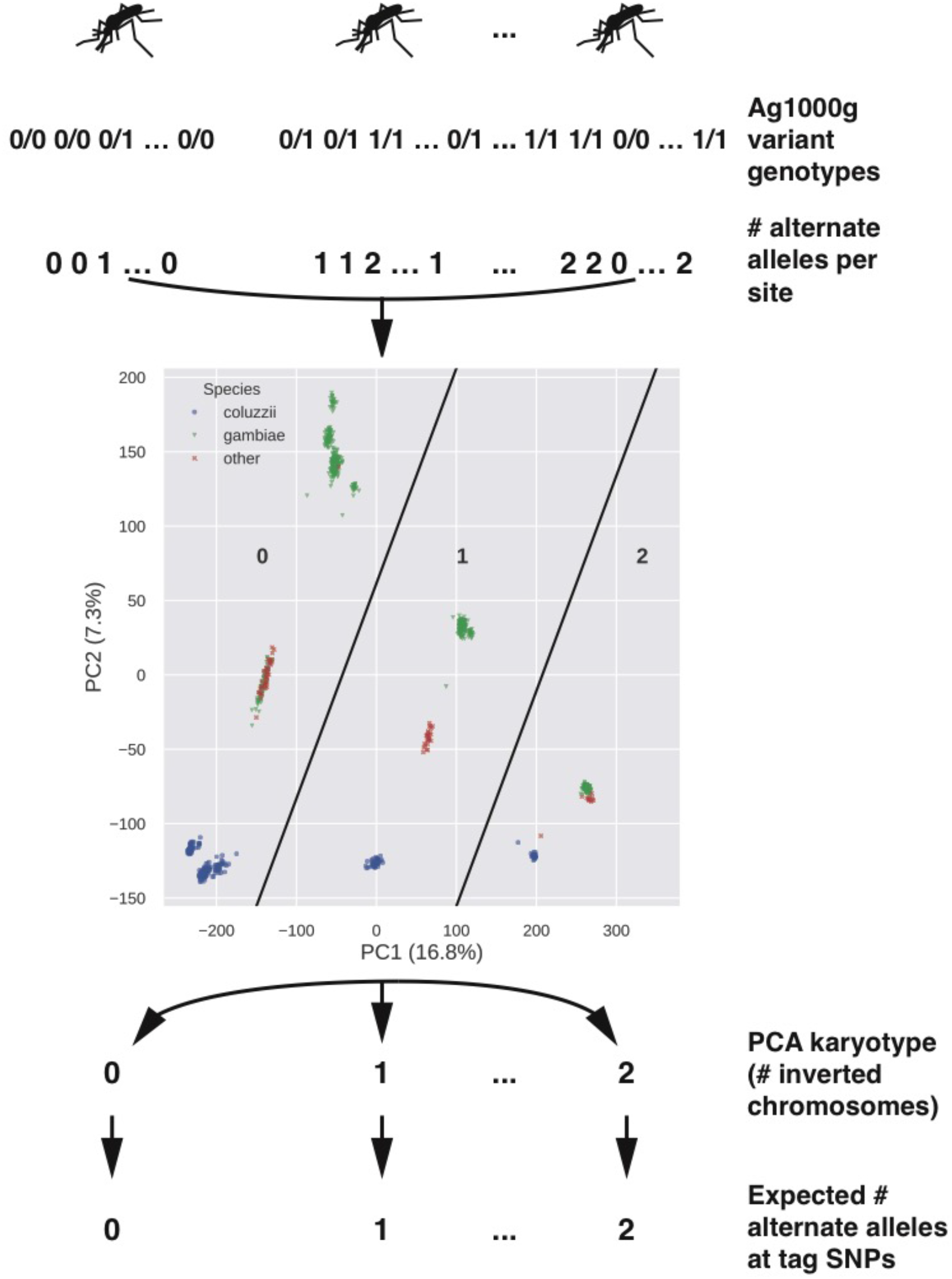
Assessment of correspondence between SNP and inversion genotype in each mosquito. For each chromosomal rearrangement and mosquito, biallelic SNP genotypes inside rearrangement boundaries were converted to a number representing the count of alternative alleles (relative to the AgamP4 reference). We applied PCA to the resulting matrix to assign each individual mosquito an inversion genotype. The expectation is that the PCA-based genotype, expressed as the number of inverted chromosomes at the focal rearrangement, should match the number of alternative alleles at SNPs predictive of inversion status (tag SNPs).

More formally, we sought to identify candidate tag SNPs using the procedure illustrated in Figure 3 (applied separately for each inversion). Specimens assigned a PCA-based karyotype for a focal inversion were divided into a training sample used for tag SNP discovery (75%) and a validation sample that was held in reserve until a later time (25%), using the model_selection module of the scikit-learn Python package (v0.19.2) (Pedregosa *et al*. 2011). We ensured that both partitions were balanced with regard to inversion genotypes but randomized in all other respects. For robust identification of candidate tag SNPs within the training sample, we masked all SNP genotypes inside the inversion boundaries with a genotype quality (GQ) below 20.

**Figure 3.**
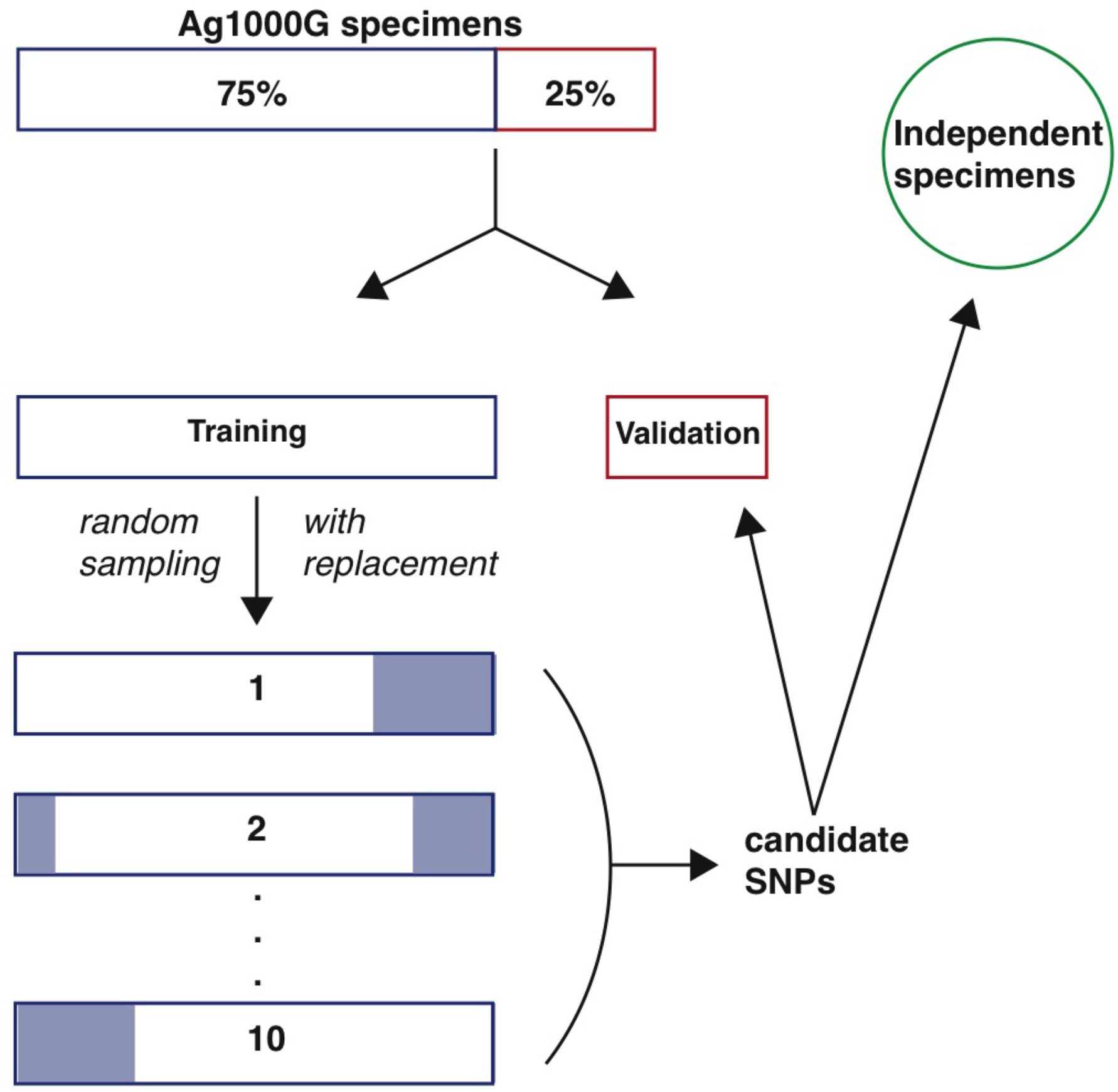
Overview of experimental design. For each inversion, the appropriate Ag1000G sample of mosquitoes that had been successfully karyotyped by PCA was partitioned into a training set (75%) and a validation set (25%). Ten bootstrap replicates of the training set were created from a random sample of 75% of the full training set. For each bootstrap replicate and each mosquito, higher frequency biallelic SNPs within inversion breakpoints were interrogated for genotypic concordance with the PCA-based genotype. Results were summarized across the ten replicates to create a set of candidate tag SNPs with concordance rates exceeding 80%. Candidate tags were used to genotype the held-out validation set, and the computational karyotype score computed across tags was compared to the PCA-based karyotype. Candidate tags were also used to interrogate mosquitoes sequenced independently of Ag1000G, and the computational karyotype score was compared to the associated cytogenetically determined karyotype.

Next, we created ten bootstrap replicates of the training sample (Figure 3). Each of the ten replicates consisted of sub-samples of 75% of the full training sample, chosen at random with respect to all variables except inversion genotype balance. For each bootstrap replicate at each interrogated SNP (biallelic, MAF≥5%), we calculated the SNP genotype-inversion genotype concordance for each mosquito in the sample, as described above (Figure 2). Genotypic concordance at each SNP interrogated in a given bootstrap replicate was expressed as the percentage of mosquitoes for which the number of alternate SNP alleles matched the number of inverted chromosomes. Because an imbalance among inversion genotypes could lead to false-positive tag SNPs, we calculated concordance separately for the three inversion genotypes in each of the ten bootstrap replicates. We then averaged the concordance scores across the ten replicates, by inversion genotype. To generate a single, conservative tag SNP concordance statistic, we used the minimum of the three mean values. Note that because the mosquito composition differed among bootstrap replicates, some SNPs were not evaluated in all ten, if they did not pass our filters in one or more iterations. Finally, to eliminate SNP positions with high levels of missing genotypes, we also calculated for each inversion genotype in each bootstrap replicate the percentage of mosquitoes with SNP genotype calls at the candidate tag (the ‘call rate’), and averaged across the ten replicates.

The procedure just described returned from 99 to 349 candidate tag SNPs for five inversions, but only two for 2R*c* (Table 1). We therefore adopted a modified approach to control for suspected population structure. One possible source of structure was the haplotype configuration of 2R*c* with respect to the flanking inversions (2R*b* and 2R*u*) (Figure 1). The inverted orientation of 2R*c* is in almost perfect linkage disequilibrium with the inverted orientation of either 2R*b* (as haplotype ‘2R*bc*’) or 2R*u* (as haplotype ‘2R*cu*’). In a ~50-year cytogenetic database compiled from samples collected in many parts of sub-Saharan Africa (described in Pombi *et al*. 2008), only four specimens were ever recorded as carrying the inverted orientation of 2R*c* unaccompanied by either 2R*b* or 2R*u* (V. Petrarca, unpublished data). A second source, not mutually exclusive, was population structure between *An. coluzzii, An. gambiae*, and the BAMAKO chromosomal form that is subsumed taxonomically within *An. gambiae* but is at least partially reproductively isolated and genetically differentiated (Manoukis *et al*. 2008; Love *et al*. 2016). Although 2R*c* occurs in all three taxa, there is a strong karyotype imbalance among them in natural populations and in Ag1000G. For example, of 70 *An. coluzzii* with 2R*c* in Ag1000G, at least 49 (70%) carried the 2R*bc* haplotype (haplotypes of the other specimens could not be inferred unambiguously). Similarly, of 64 non-BAMAKO *An. gambiae* with 2R*c*, 62 (97%) carried the 2R*bc* haplotype. On the other hand, all 45 BAMAKO, by definition, carried 2R*cu*. We initially partitioned our sample by species, but the inclusion of BAMAKO in the *An. gambiae* partition resulted in very few candidate tags concordant with inversion genotype (N=17). Ultimately, we retained two data partitions (*An. coluzzii* and non-BAMAKO *An. gambiae*), eliminating a third BAMAKO partition due to the fixation of 2R*c* in this taxon (Coluzzi *et al*. 1985). From the non-BAMAKO *An. gambiae* partition (hereafter, *An. gambiae* for brevity), we omitted two of only three specimens carrying 2R*u* (AZ0267-C from Mali and AV0043 C from Guinea), guided by PCA. As described above, both data partitions were split into training (75%) and validation (25%) sets, and ten bootstrap replicates of each training set were analyzed.

**Table 1.**
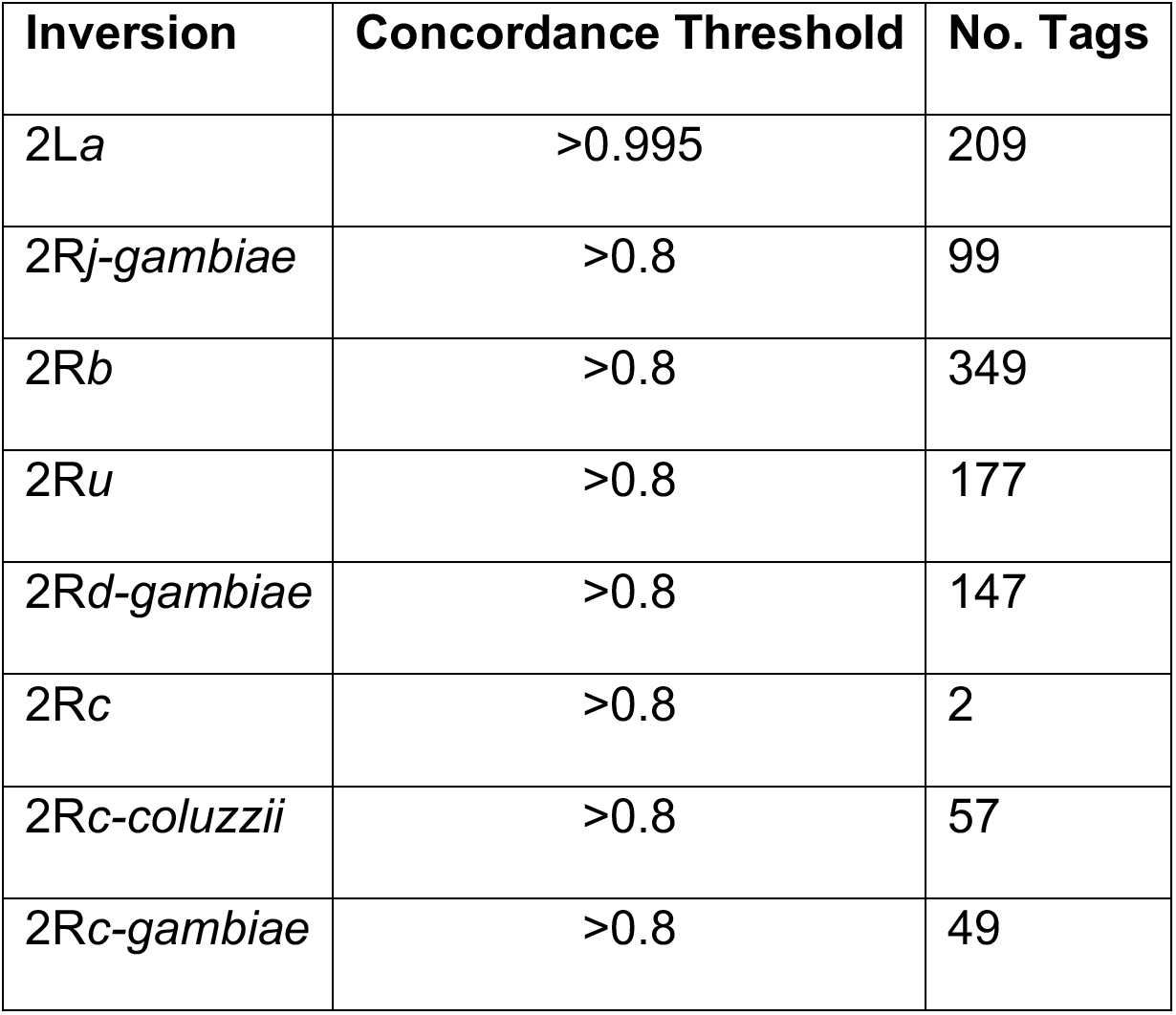
Candidate tag SNPs predictive of inversion genotype in Ag1000G data

| Inversion | Concordance Threshold | No. Tags |
| --- | --- | --- |
| 2La | >0.995 | 209 |
| 2Rj-gambiae | >0.8 | 99 |
| 2Rb | >0.8 | 349 |
| 2Ru | >0.8 | 177 |
| 2Rd-gambiae | >0.8 | 147 |
| 2Rc | >0.8 | 2 |
| 2Rc-coluzzii | >0.8 | 57 |
| 2Rc-gambiae | >0.8 | 49 |

Ultimately, the candidate tag SNPs chosen (Table 1) met the following three criteria: they were (i) analyzed in at least eight of the ten bootstrap replicates; (ii) called at a rate greater than 90% within each karyotype class; and (iii) concordant with karyotype more than 80% of the time within each karyotype class (99.5% for 2L*a*). Their approximate physical position relative to the span of each inversion is illustrated in Figure S1.

### Validation of candidate tag SNPs in Ag1000G

We interrogated the candidate tag SNPs in the validation samples from Ag1000G that had been held aside during the discovery phase (Figure 3). For each mosquito in the validation set, we masked genotypes inside the focal inversion with GQ scores less than 20. Next, among the retained SNPs, we identified those corresponding to candidate tags and converted their diploid genotypes to a count of the number of alternate alleles. Finally, the number of alternate alleles at each tag SNP was summed across tags and averaged to provide an overall computational karyotype score. We compared this mean score to the PCA-based karyotype.

### Testing tag SNPs in data independent of the Ag1000G pipeline

We also explored the efficacy of our tag SNPs for computational karyotyping in wild-caught mosquitoes subject to whole genome sequencing and variant calling by individual investigators, for which corresponding cytological karyotypes had been determined through phase microscopy (Figure 3). We used specimens originating from southern Mali, 8 *An. gambiae* BAMAKO chromosomal form (Fontaine *et al*. 2015; Love *et al*. 2016) and 17 *An. coluzzii* (Main *et al*. 2015), whose variant calls and cytogenetic metadata are publicly accessible (Table S5). These data include specimens sequenced to much lower coverage than the standard adhered to by Ag1000G. We followed the same procedure described for the Ag1000G validation set to computationally karyotype these specimens, and compared their computational and cytologically determined karyotypes.

### Genetic distance trees to assess inversion history

We compared patterns of relatedness near the breakpoints of all six inversions using unrooted neighbor-joining (NJ) trees. For each inversion, we used biallelic SNPs with a MAF of 0.01 found within 5 kb upstream and downstream of the distal and proximal breakpoints (15 kb for 2R*d*). Total numbers of SNPs for each inversion were: 2L*a*, 596; 2R*j*, 909; 2R*b*, 428; 2R*c*, 2141; 2R*d*, 955; 2R*u*, 1110. Using the python package *anhima*, we converted the number of alternate alleles at these SNPs into a Euclidean distance matrix, and then constructed neighbor-joining trees using all 1,347 specimens. To assess support for the nodes of the 2R*c* tree, we used the transfer bootstrap estimate (TBE; Lemoine *et al*. 2018), a statistic that measures the number of taxa that must be transferred to make a given branch of a reference tree match the closest equivalent branch in a bootstrap tree. To calculate this statistic, we imported the matrix of alternate allele counts into R (v. 3.5.1, “Feather Spray”; R Core Team 2018) and used the dist() function of base R to construct the Euclidean distance matrix. We then used the nj() function in the ape package (v. 5.2) to construct the neighbor joining tree, and the boot.phylo() function to generate 1,000 bootstrap trees. We used these trees as input to booster (Lemoine *et al*. 2018), which calculates the TBE for each node.

### Code and data availability

All genomic sequence data and variant call files used in this study are located in open data repositories as specified in Tables S1 and S2. The *An. gambiae* AgamP4 reference assembly is available through VectorBase (https://www.vectorbase.org/). All custom code necessary to reproduce this analysis can be found at https://github.com/rrlove/comp_karyo_notebooks and https://github.com/rrlove/ingenos. The complete set of tag SNPs, together with a custom script for computational karyotyping, which calculates the mean inversion genotype across the relevant tag SNPs, can be found at https://github.com/rrlove/compkaryo.

## Results

After filtering, we retained the genotype data from 1,347 individually sequenced *An. coluzzii* and *An. gambiae* mosquitoes from the Ag1000G repository of natural genomic sequence variation, representing population samples from 13 West, Central, and East African countries and the island of Mayotte (Tables S1, S2).

### Patterns of genetic variation at inversion breakpoints

To gain insight into the relative roles of inversion history, taxonomic status, and geographic location in structuring genetic variation for each inversion, we reconstructed neighbor-joining trees based on SNPs in the immediate vicinity of the breakpoints (Figure 4). The resulting dendrograms, color-coded by inversion genotype, taxon and African country, indicate little clustering on the basis of geographic location; outlier population samples are those with a history of inbreeding or hybridization (see Methods). On the other hand, with the notable exception of 2L*a*, taxonomic status is an important factor structuring inversion variation between *An. gambiae* and *An. coluzzii*. Moreover, BAMAKO specimens appear to comprise a differentiated outlier clade within the larger *An. gambiae* cluster. It is interesting to note that for inversion 2R*c*, taxonomic status appears to be a more decisive factor than inversion genotype. All three 2R*c* inversion genotypes cluster within their respective species (supported by bootstrap at 90%, or 98% if dendrograms are constructed after removing outlier samples from The Gambia, Guinea-Bissau and Kenya; not shown). Further investigation is required to determine whether this pattern results from a monophyletic inversion that subsequently differentiated between taxa, or from independent inversion events in the two taxa.

**Figure 4.**
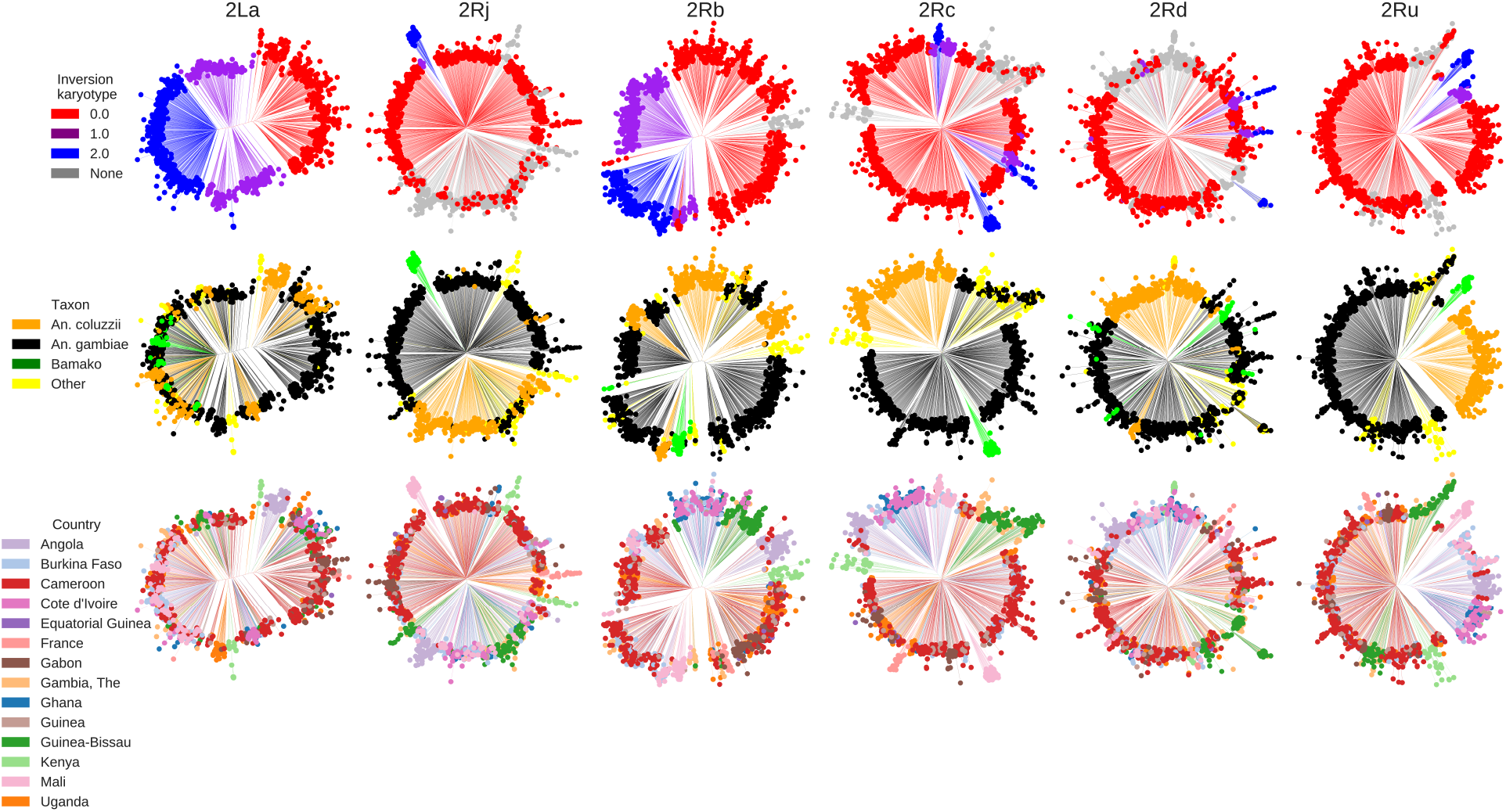
Neighbor-joining dendrograms reconstructed from 1,347 *An. gambiae* and *An. coluzzii* mosquitoes from Ag1000G, using biallelic SNPs within 5 kb of inversion breakpoints (15 kb for 2R*d*) having a minimum minor allele frequency of 0.01. Columns represent the same inversion dendrogram, alternately color-coded by inversion genotype as determined from PCA (first row), taxon (second row), or geographic source (third row). Some specimens that could not be karyotyped by PCA for inversions 2R*c*, 2R*d*, 2R*j*, and 2R*u* had cytogenetically determined karyotypes, which were used in place of PCA for color-coding the inversion genotype. ‘None’ refers to mosquitoes that were not assigned an inversion genotype either by PCA or cytogenetically; ‘Other’ refers to mosquitoes that were not identified taxonomically.

### Inversion karyotype imputation by PCA

Only 373 of the 1,347 mosquitoes were associated with metadata that included cytologically determined inversion karyotypes. As discovery of candidate tag SNPs requires provisional inversion genotype assignments, we applied local PCA to assign genotypes for individual inversions on chromosome 2, following Ma and Amos (2012). A recognized limitation to this population-level approach, beyond the fact that it cannot be applied to individual mosquitoes, is that its success depends upon the presence of all three inversion genotypes in the sample under study. For this reason, and with the goal of finding the most flexible solution to inversion genotyping across geography and taxa, we began with PCA based on the entire set of 1,347 mosquitoes, under the simplifying assumption that the expected ‘three-stripe’ signal on a PCA plot would not be overwhelmed by geographic or population structure. Only in the case of 2L*a* could genotype assignments be confidently inferred from the combination of all 1,347 specimens. For inversions on 2R, from one to four admixed (*An. gambiae-An. coluzzii*) or geographic outlier populations (highly inbred or island samples) had to be removed from analysis before a three-genotype pattern could be discerned on the PCA plot (Tables S2, S4; see Methods). Additionally, for 2R*d* and 2R*j*, *An. gambiae-An. coluzzii* population structure dominated the PCA. Taken together with the fact that 2R*j* has yet to be found in *An. coluzzii* (Coluzzi *et al*. 2002; della Torre *et al*. 2005), we removed all 341 *An. coluzzii* specimens (Tables S2, S4) prior to PCA karyotyping of 2R*d* and 2R*j* in *An. gambiae*. Ultimately, PCA karyotypes were imputed for 780-1,347 mosquitoes, depending upon the inversion (Table S4).

### Tag SNP ascertainment and validation in Ag1000G

Dividing the Ag1000G samples into training (75%) and validation (25%) sets for each inversion, and working within the training sets using a bootstrapping procedure, we screened for candidate tag SNPs in the five 2R inversions and 2L*a* (see Methods for details). Candidate tag SNPs were those whose genotypes were concordant with the corresponding PCA genotypes, when averaged across ten bootstrap replicates, for more than 80% of the specimens that were scored (99.5% for 2L*a*). The number of candidate tags ranged from 99 (2R*j*) to 349 (2R*b*) excluding 2R*c*, in which only two candidates were found due to population structure between *An. gambiae* and *An. coluzzii* (Figure 4; Table 1). Partitioning the 2R*c* sample by taxon (and omitting BAMAKO; see Methods) resulted in 49 and 57 tags for *An. gambiae* and *An. coluzzii*, respectively (Table 1). Notably, there was no overlap between the two sets of tags.

To assess the performance of these candidate tags, we used them to predict karyotypes in the held-out validation sets of Ag1000G specimens. For each inversion and specimen, we calculated a computational karyotype score representing the average genotype inferred across all candidate tag SNPs ascertained (see Methods). Histograms of resulting computational karyotype scores generally showed tight clustering around the three theoretical genotypic optima (0, 1, 2), reflecting close agreement among specimens (Figure 5). For each specimen in a validation set, we then compared the computational karyotype score to its PCA karyotype, and tallied the number of disagreements (Table 2). All except one specimen had matching PCA and computational karyotype scores. This exception, one of 254 (0.4%) assignments for 2R*c* in *An. gambiae*, involved a specimen carrying 2R*u* (AZ0267-C) already noted as an outlier (see Methods).

**Figure 5.**
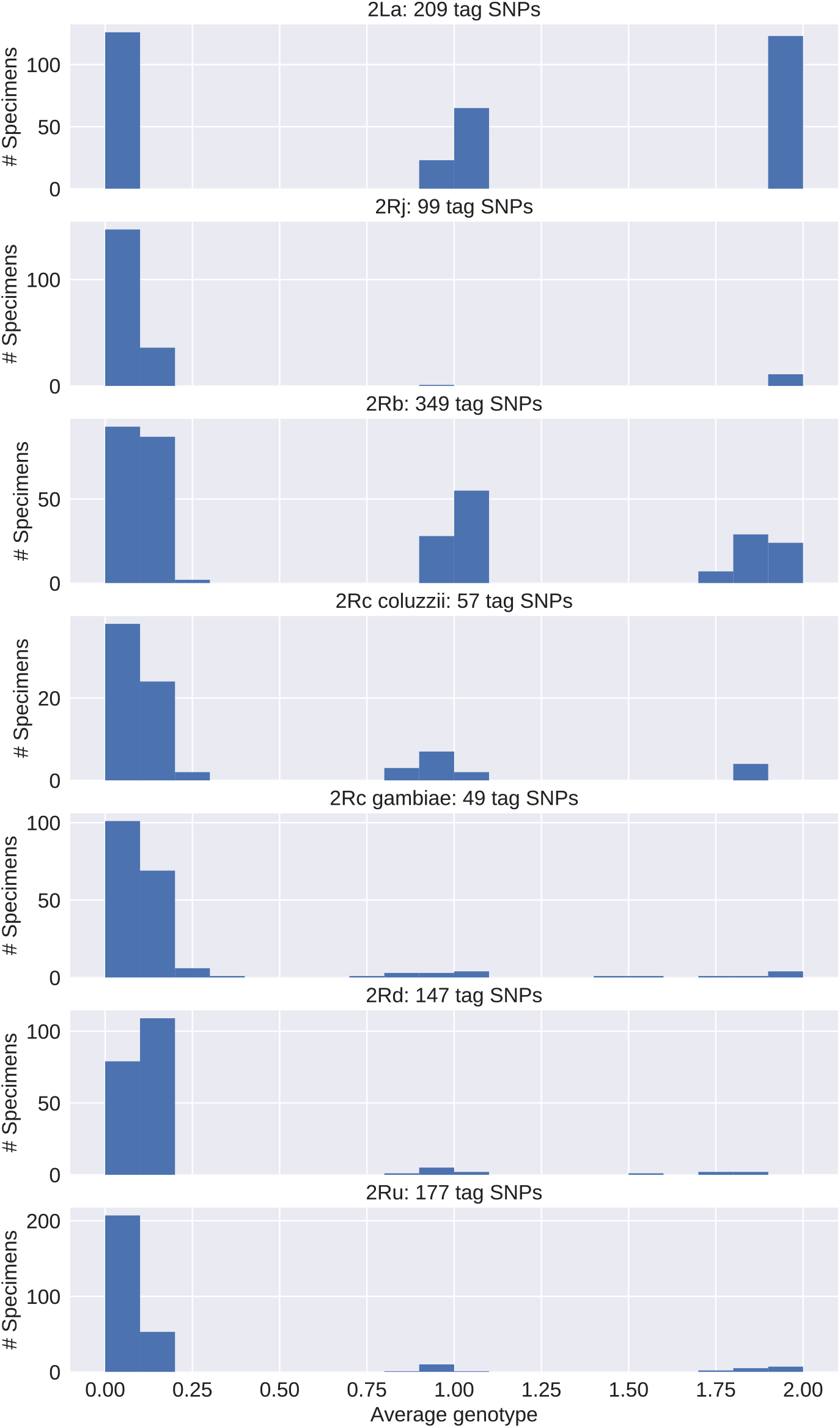
Histograms of computational karyotyping scores calculated by interrogating tag SNPs in *An. gambiae* and *An. coluzzii* mosquitoes from the Ag1000G validation sets. Note that these mean scores cluster around 0, 1, and 2.

**Table 2.** Mismatches between PCA and computational karyotypes in the Ag1000G validation sets

| **Inversion** | **Total specimens** | **No. tags scored** | **Matching karyotypes** |  | **Mismatched karyotypes** |  |
| --- | --- | --- | --- | --- | --- | --- |
|  |  |  | **No. specimens** | **% tags supporting score** | **No. Specimens** | **% tags supporting score** |
| 2La | 337 | 168-203 | 337 | 93.6-100 | 0 | -- |
| 2Rj-<br>*gambiae* | 195 | 94-99 | 195 | 83.8-100 | 0 | -- |
| 2Rb | 325 | 304-349 | 325 | 77.5-97.7 | 0 | -- |
| 2Rc-<br>*coluzzii* | 80 | 55-57 | 80 | 78.9-100 | 0 | -- |
| 2Rc-<br>*gambiae* | 196 | 45-49 | 195 | 59.6-100 | 1 | 67.3 |
| 2Rd-<br>*gambiae* | 201 | 128-147 | 201 | 55.2<sup>1</sup>-95.9 | 0 | -- |
| 2Ru | 286 | 124-177 | 286 | 76.6-100 | 0 | -- |

<sup>1</sup>Next highest value is 70.1%.

### Performance of tag SNPs in resequencing data independent of Ag1000G

Previous studies re-sequenced *An. gambiae* or *An. coluzzii* mosquitoes from Mali whose karyotypes had been determined from the polytene chromosome banding pattern (Main *et al*. 2015; Love *et al*. 2016). Although sample size is limited, these data allow a direct comparison of cytogenetic and *in silico* karyotyping under less ideal conditions—lower sequencing depth, with variant calls made independently of the Ag1000G pipeline. For each specimen and inversion, we calculated computational karyotype scores (averaged across all tag SNPs that could be ascertained in a specimen) (Tables S5, S6). Histograms of these scores by inversion, similar to those based on Ag1000G validation sets, reveal clustering of scores around the three genotypic optima provided that taxon-specific tags (2R*c-coluzzii* and 2R*c-gambiae*) are applied to the conspecific taxon, and heterospecific applications (including use of 2R*c-gambiae* tags to genotype BAMAKO) are avoided (Figure 6, Figure S2).

**Figure 6.**
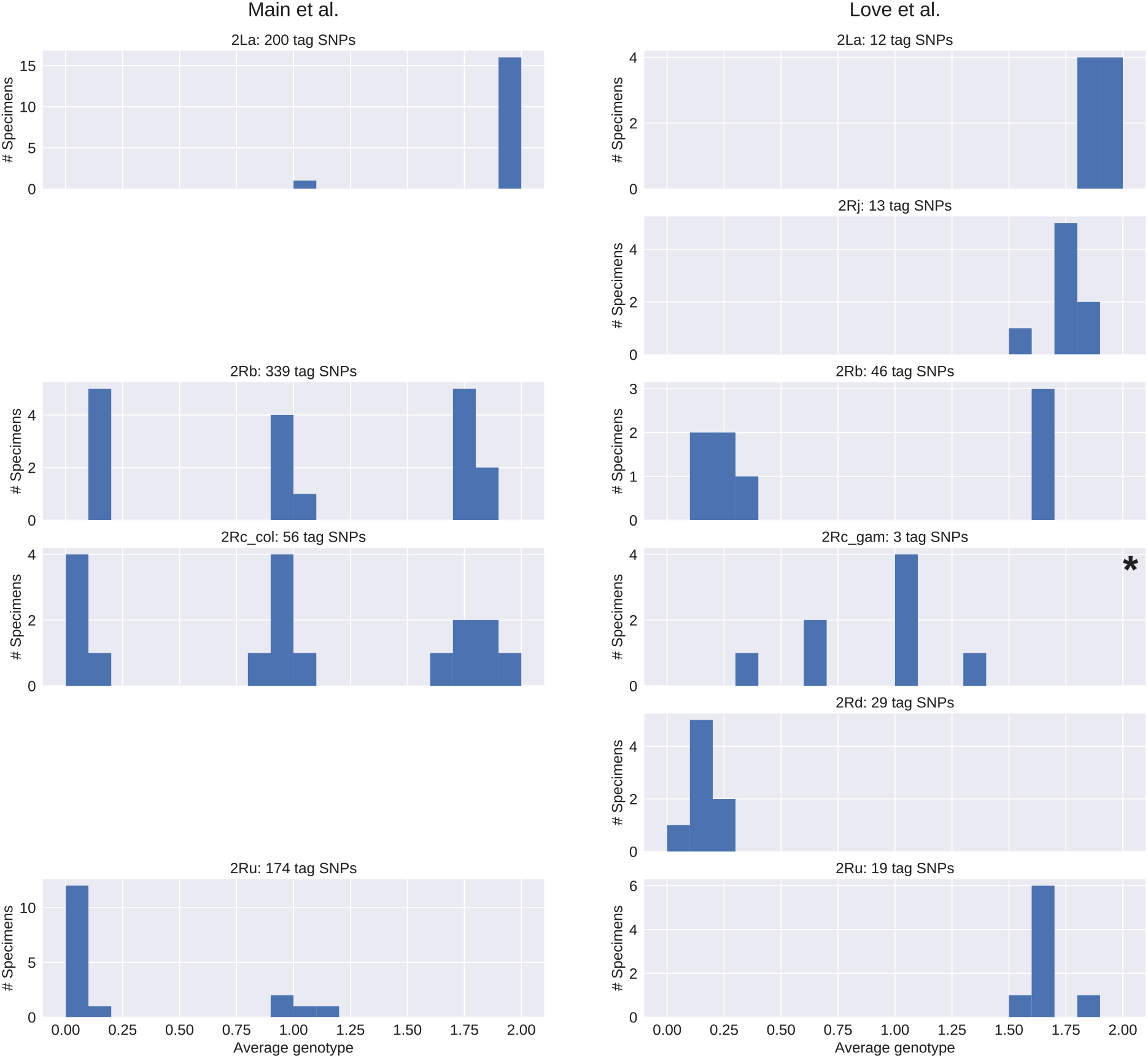
Histograms of computational karyotyping scores calculated by interrogating tag SNPs in *An. gambiae* and *An. coluzzii* mosquitoes re-sequenced independently of the Ag1000G pipeline, often at lower sequencing depth. Scores cluster near 0, 1, and 2 with little dispersion except when taxon-specific tag SNPs are applied to a different taxon (indicated by an asterisk).

In the BAMAKO sample of Love *et al*. (2016) where mean sequencing depth ranged from 9-10x, there was concordance in karyotype assignments between cytogenetic and computational methods for five inversions including 2L*a*, even though only 10-12 2L*a* tags were ascertained (Table S5, S6). However, as expected for BAMAKO, the *An. gambiae* 2R*c* tags failed. Due to the extreme geospatial restriction of BAMAKO, this specific problem is limited in scope.

In the *An. coluzzii* sample of Main *et al*. (2015), mean sequencing coverage varied widely (4-66x; Tables S5, S6). The impact of very low sequencing coverage on the success of computational karyotyping is illustrated by specimens 04SEL021 and 04SEL02 (4x and 5x, respectively). For 04SEL021, there is no apparent disagreement between the cytogenetic and mean computational genotype scores for any of the six inversions. Nevertheless, for those inversions classified as heterozygotes both cytogenetically and computationally (2L*a*, 2R*b*, 2R*c*), the proportion of tags whose genotype matches the mean computational score drops drastically to ~30% (Table S5), likely because true heterozygous sites are often scored as homozygous either for the reference or alternate allele (0 or 2) due to low sequencing coverage. (Indeed, using chromosome 3L, we confirmed the expected drop in the rate of heterozygosity with decreasing coverage in these 17 specimens; data not shown). Low coverage alone is less likely to bias computational scores toward zero or two. For 04SEL02 (5x coverage), where cytogenetic versus computational discrepancies occur at 2R*b* and 2R*u*, the computational karyotype is supported respectively by 81% of 208 tags and >94% of 57 tags, favoring the computational genotype by weight of evidence. The remaining six specimens with discordant inversion genotypes were sequenced to at least 10x coverage. In these cases, when the computational genotype score signaled ‘1’ in contradiction to a homokaryotypic cytogenetic genotype (02SEL85, 02SEL006, 02SEL009, 01Osel134), the proportion of tags supporting the computational genotype ranged from 65% to >92%. For other types of genotypic disagreements between methods, the computational score was supported by >80% of tags scored. Overall, these results suggest that computational karyotyping using tag SNPs can be successful in data derived independently of Ag1000G (Tables S5, S6), though care should be taken when this approach is applied to very low coverage samples.

### Performance of tag SNPs against cytogenetically karyotyped Ag1000G specimens

We compared the cytogenetic karyotype assignments for 373 specimens in Ag1000G to their corresponding computational karyotype assignments (Table 3). Conflicts were few overall, and for every inversion, all but one conflict (involving specimen AZ0267-C, the exceptional *An. gambiae* carrier of the 2R*u* inversion) could be attributed to errors in the cytogenetically assigned scores, as genotypes imputed from both PCA and tag SNPs contradict the cytogenetic assignment. Visual reference back to the PCA plots clearly confirmed that for specimens whose cytogenetic and tag SNP assignments differed and for whom PCA karyotypes could be determined, their locations on the plot strongly agreed with the tag SNP genotype (Figure S3). Considering that we ascertained tens or hundreds of tags per specimen, and that the proportion of tags whose SNP genotype matched the computational score was greater than 83% in all except the unusual specimen AZ0267-C (Table 3), the computational scores more confidently predict the true inversion genotype than traditional cytogenetics for these five inversions. The most dramatic example is with respect to inversion 2R*u*, where we noted an unusually large number of erroneous cytogenetic genotypes of ‘1’ (N=18/29) conflicting with both PCA and computational assignments of ‘0’. It is not immediately clear what could lead to such an elevated rate of cytogenetic error (which otherwise is ~4%), but it is possible that the 2R*u* heterozygous loop was mistaken either for a loop created by a rare inversion (sensu Pombi *et al*. 2008) in the same chromosomal region, or for a 2R*d* loop in samples from regions where 2R*u* is rare (as supported by the fact all 11 cytogenetic errors in *An. gambiae* were found in samples from the same small region in Cameroon, six of which were scored computationally as ‘1’ for 2R*d*).

**Table 3.** Discrepancies between cytogenetic and computational karyotypes in Ag1000G mosquitoes analyzed.

| Inversion<br>Tags | Partition | CYT | Specimens<br>(N) | Specimens with discrepancies |  |  |
| --- | --- | --- | --- | --- | --- | --- |
|  |  |  |  | Mismatch CYT-TAG<br>(%) | Match TAG-PCA<br>(%) | No. tag SNPs<br>scored<br>(% matching TAG) |
| **2La** |  | 0 | 117 | 5 (4.3) | 5 (100) | 200-203 (99.5-100) |
|  |  | 1 | 68 | 5 (7.4) | 5 (100) | 193-203 (100) |
|  |  | 2 | 160 | 2 (1.3) | 2 (100) | 201-203 (99.5-100) |
| **2Rj-gambiae** | *gambiae* | 0 | 236 | 0 (0) | -- | — |
|  |  | 1 | 4 | 0 (0) | -- | — |
|  |  | 2 | 45 | 0 (0) | -- | — |
| **2Rb** |  | 0 | 127 | 2 (1.6) | 2 (100) | 348 (85.3-87.6) |
|  |  | 1 | 124 | 4 (3.2) | 4 (100) | 346-349 (86.8-93.7) |
|  |  | 2 | 121 | 6 (5.0) | 6 (100) | 331-348 (88.8-91.6) |
| **2Rc-gambiae** | *gambiae*<sup>1</sup> | 0 | 184 | 7 (3.8) | 7 (100) | 48-49 (83.7-98.0) |
|  |  | 1 | 32 | 3 (9.4) | 2 (66.7) | 47-49 (42.9<sup>2</sup>-91.5) |
|  |  | 2 | 24 | 2 (8.3) | 2 (100) | 48-49 (90.0-91.8) |
| **2Rc-coluzzi** | *coluzzii* | 0 | 13 | 1 (7.7) | 1 (100) | 56 (87.5) |
|  |  | 1 | 25 | 0 (0) | -- | — |
|  |  | 2 | 16 | 0 (0) | -- | — |
| **2Rd-gambiae** | *gambiae* | 0 | 234 | 9 (3.8) | 9 (100) | 143-147 (84.9-96.6) |
|  |  | 1 | 28 | 4 (14.3) | 4 (100) | 146-147 (88.4-93.9) |
|  |  | 2 | 22 | 3 (13.6) | 3 (100) | 146-147 (89.1-91.8) |
| **2Ru** | *col+gam* | 0 | 263 | 1 (0.38) | 1 (100) | 176 (85.2) |
|  |  | 1 | 29 | 18 (62.1) | 18 (100) | 170-177 (88.7-99.4) |
|  |  | 2 | 47 | 1 (2.1) | 1 (100) | 176 (97.2) |

CYT, cytogenetic genotype; TAG, computational genotype; PCA, genotype inferred by PCA.
<sup>1</sup>*An. gambiae* excluding BAMAKO
<sup>2</sup>This value corresponds to one of three non-BAMAKO *An. gambiae* carriers of the 2Ru inversion, AZ0267-C. The next highest value is 85.7.

Our results also highlight the pitfalls of using taxon-specific tags to genotype other taxa, or populations with high levels of admixture between taxa (Table S7). As expected, we find elevated numbers of cytogenetic-computational disagreements when (i) 2R*c-gambiae* tags are applied to BAMAKO (60% of the 45 specimens), (ii) 2R*d-gambiae* tags are used to genotype *An. coluzzii*, and (iii) 2R*d-gambiae* tags are applied to admixed *An. gambiae-An. coluzzii* populations such as those from Guinea Bissau. These disagreements involve specimens carrying inverted arrangements according to cytogenetic analysis which are not tagged as inverted computationally, due to the lack of correlation between tags and the inverted orientation in the heterospecific genetic background.

## Discussion

Analysis of the Ag1000G database allowed us to develop the first standardized computational karyotyping of the six main polymorphic chromosomal inversions in the major malaria vectors *An. coluzzii* and *An. gambiae*, despite the fact that only a small subset of specimens in the database had cytogenetic karyotype assignments (Figure 7). Direct comparison of computational karyotype scores with the cytogenetic assignment for the same specimen in Ag1000G suggests that computational karyotyping outperforms traditional cytogenetics in terms of accuracy, given that assignments are based on tens or hundreds of individual tags. Preliminary testing on specimens sequenced and computationally processed by individual laboratories outside of Ag1000G standards suggests that our tag SNPs have the potential to perform well, even on specimens sequenced to much lower depth. Our approach not only has a lower error rate compared to classical cytogenetics, but also is more widely applicable (regardless of mosquito gender, physiological status, or method of preservation), more widely accessible to those without specialized expertise, higher throughput, and therefore, ultimately cheaper to implement at scale. This method can now be used to predict inversion genotypes in previously sequenced data sets for which ecological and behavioral data may already be available. Even more important, easy large-scale adoption of this approach allows for new and properly powered association studies to be performed on ecologically and epidemiologically relevant mosquito phenotypes, studies that that would have been prohibitively ambitious when relying on cytogenetic karyotyping. In addition, this method can now facilitate sequencing experiments for which inversion karyotype is relevant at scales. Expanding the possibilities further, molecular assays based on these results that will allow inversion genotyping without whole genome sequencing are under development.

**Figure 7.**
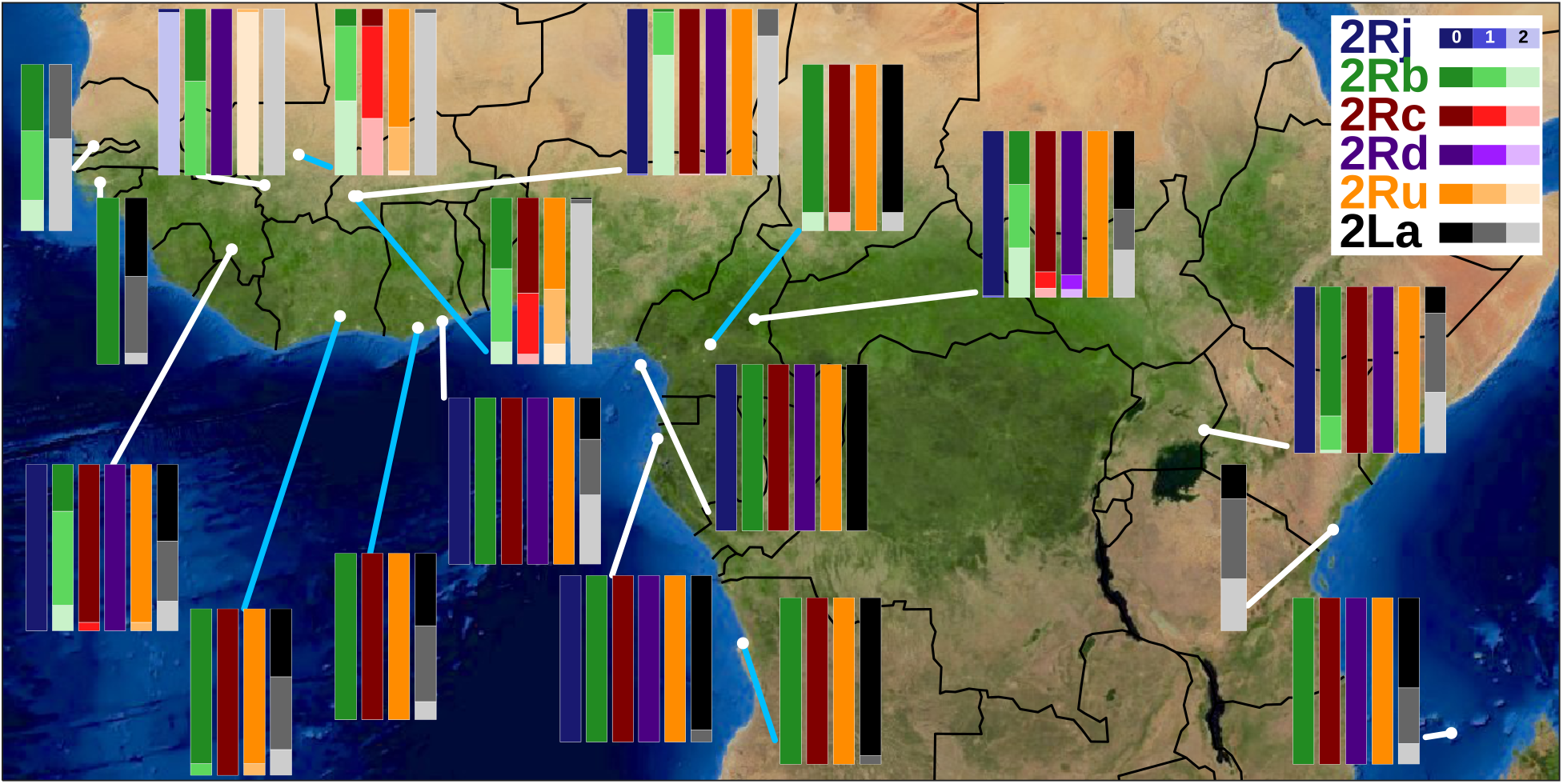
Map of the study area with the frequency of *An. gambiae* and *An. coluzzii* inversion genotypes inferred for up to six polymorphic chromosome 2 inversions summarized by country (and the island of Mayotte; see Table S2). Blue connecting lines point to *An. coluzzii* samples, while white connecting lines point to *An. gambiae* and hybrid/outlier populations. Image: Visible Earth, NASA. Produced with cartopy v.0.17.1.

However, some important limitations exist. Computational karyotyping is strictly dependent upon tag SNPs that are strongly correlated with inversion status, a contingency that depends upon representative sampling. Although Ag1000G is populated by samples derived from multiple countries in West, Central and East Africa, *An. coluzzii* is underrepresented, as is southern Africa (Miles *et al*. 2017). Even more importantly, with the exception of the cosmopolitan inversions 2L*a* and 2R*b*, the inverted orientation of other rearrangements (2R*j*, 2R*c*, 2R*d*, and 2R*u*) is underrepresented in the Ag1000G data that was available at the time of our analysis. It is clear that population structure is an especially important factor in the application and further development of tags for 2R*c* and 2R*d*. The current taxon-specific tags should not be used to genotype heterospecific specimens (including BAMAKO) or samples from areas where high rates of interspecific hybridization are known. The presence of strong population structure means that correlations between tags and the inverted orientation characteristic of the target taxon cannot be assumed in a different taxon. The absence of correlation should downwardly bias the computational score, resulting in false negatives when genotyping true inverted homozygotes and heterozygotes. Finally, our inversion breakpoint dendrograms raise the possibility that at least one cytologically-recognized inversion, 2R*c*, may have arisen repeatedly at the molecular level, a result that requires further investigation beyond the scope of this study. With the exception of 2R*c*, 2R*d*, and 2R*j*, for which we developed taxon-specific tags, our approach implicitly assumed that inversions shared by *An. gambiae* and *An. coluzzii* are monophyletic, and may yield unexpected results if this assumption is violated. Accordingly, these tools should be applied with caution, and there is ample room for improvement as more data become available. Fortunately, our standardized approach makes it easy to accommodate improvements. The success of our method thus far suggests that the general approach may be suitable for studying inversions more broadly, in additional malaria vectors as well as other systems where inversions are implicated in local adaptation.

Nearly twenty years ago, Coluzzi and colleagues predicted that the then-newly-assembled *An. gambiae* reference genome would facilitate our analyses of polymorphic chromosomal inversions in the *An. gambiae* complex (Coluzzi *et al*. 2002). Our work continues the realization of that prediction by providing, for the first time, cross-continent diagnostics for multiple inversions. These computational diagnostics, and the molecular diagnostics that they leverage, take us one step closer to understanding the contribution of chromosomal inversions to the deadly facility of *An. gambiae* and *An. coluzzii* for vectoring malaria.

## Acknowledgements

We thank the Notre Dame Center for Research Computing for technical support, and C. Liu, C. Sweet, and J. Young for helpful discussions. We thank M. Kern and R. Montanez-Gonzalez for assistance with DNA extraction. This work was supported by the National Institutes of Health (R01 AI125360 awarded to NJB). During this work, NJB was supported by Target Malaria, which receives core funding from the Bill & Melinda Gates Foundation and from the Open Philanthropy Project Fund, an advised fund of Silicon Valley Community Foundation.

## Notes

https://figshare.com/projects/Data_for_In_silico_karyotyping_of_chromosomally_polymorphic_malaria_mosquitoes_in_the_Anopheles_gambiae_complex_/65522

