## Supplemental figures and tables for "In silico karyotyping of chromosomally polymorphic malaria mosquitoes in the *Anopheles gambiae* complex"

### Table of Contents:

| | |
|---|---|
| **Figure S1** | Page 2 |
| **Figure S2** | Page 3 |
| **Figure S3** | Page 4 |
| **Table S1** | Page 5 |
| **Table S2** | Page 10 |
| **Table S3** | Page 11 |
| **Table S4** | Page 12 |
| **Table S5** | Page 13 |
| **Table S6** | Page 14 |
| **Table S7** | Page 15 |
| **Table S8** | Page 16 |

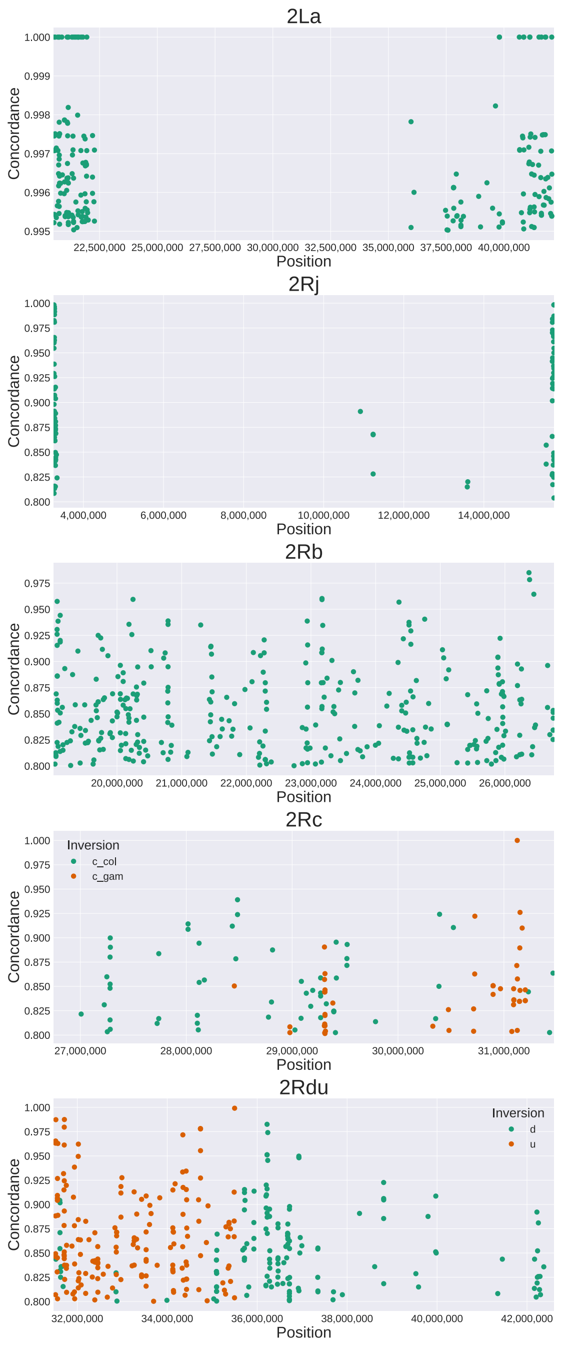

**Figure S1. Position of tag SNPs within each chromosomal rearrangement.** Scatter plots of genomic location and SNP genotype-inversion genotype concordance for tag SNPs identified for each of six inversions (2R*d* and 2R*u*, which overlap, are shown on the same plot in different colors).

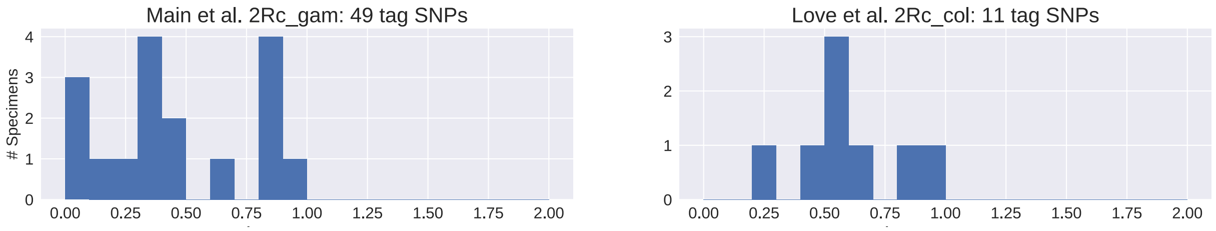

**Figure S2. Heterospecific application of taxon-specific tags results in average genotype scores that do not cluster near 0, 1, or 2**. Shown are applications of 2R*c*-*gambiae* tags to (left), the *An. coluzzii* specimens of Main *et al.* (2015) and (right), the BAMAKO specimens of Love *et al.* (2016)

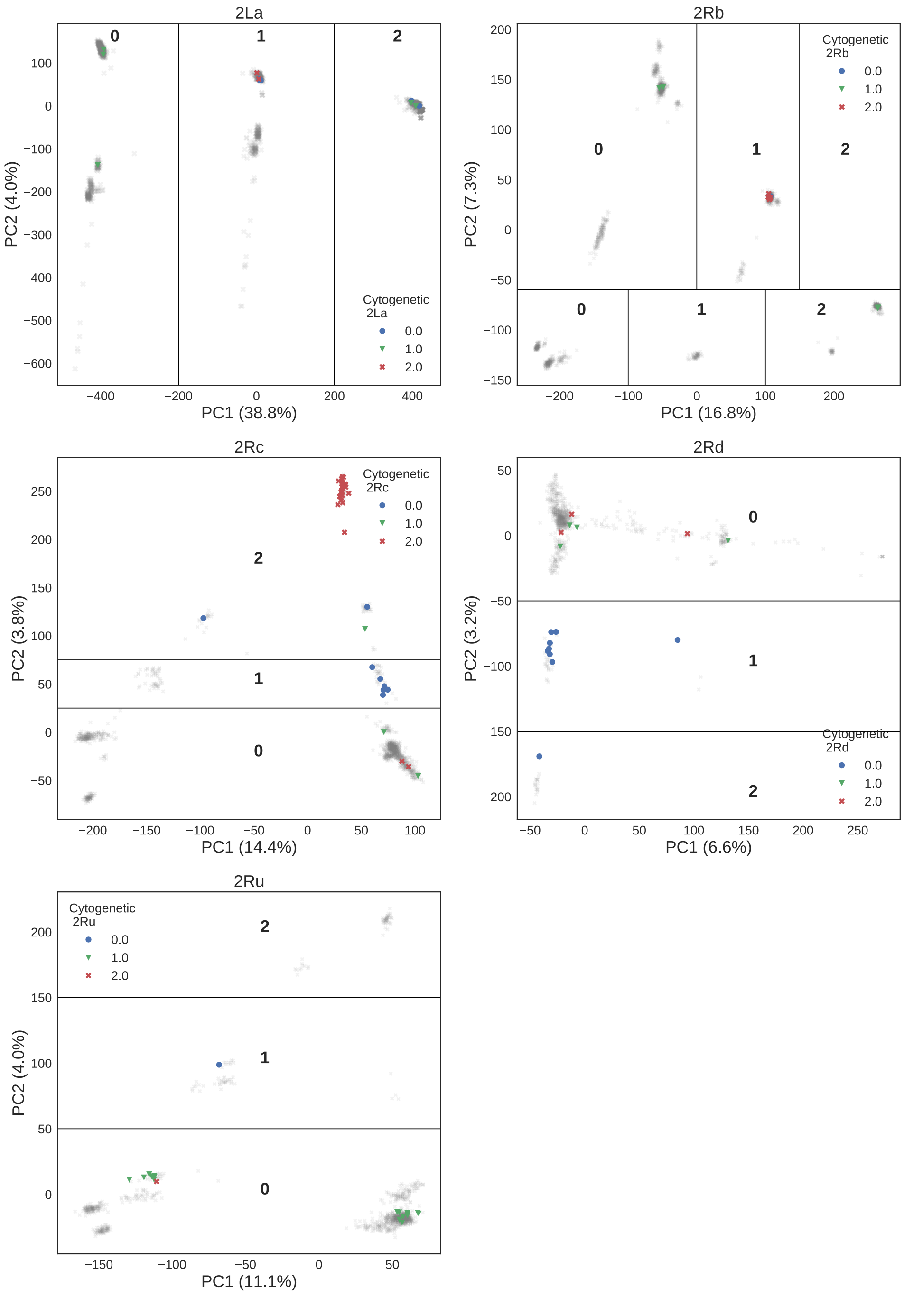

**Figure S3. Mismatches between traditional cytogenetics and computational assignments are likely cytogenetic errors.** Shown are PCA plots based on genomic windows covering each rearrangement, in which colored dots represent individual specimens from Table 3 whose cytogenetic assignment disagreed with PCA and computational karyotype assignments. Partitions are labeled by karyotype. Upper and lower partitions in the 2R*b* PCA plot correspond to *An. gambiae* and *An. coluzzii*, respectively.

**Table S1.** Specimen IDs and ENA accessions for Ag1000G phase 3 and MalariaGEN Vector Observatory *An. gambiae* and *An. coluzzii* mosquitoes not yet publicly released by MalariaGEN.

| ENA | Sanger_ID | Ox_code | Country | Species |
|---|---|---|---|---|
| ERS1119029 | 2572STDY6370039 | AN0477-C | Cameroon | *An. gambiae* |
| ERS1119030 | 2572STDY6370040 | AN0478-C | Cameroon | *An. gambiae* |
| ERS1119009 | 2572STDY6370019 | AN0444-C | Cameroon | *An. gambiae* |
| ERS1119010 | 2572STDY6370020 | AN0445-C | Cameroon | *An. gambiae* |
| ERS1118989 | 2572STDY6369995 | AN0418-C | Cameroon | *An. gambiae* |
| ERS1118990 | 2572STDY6369996 | AN0419-C | Cameroon | *An. gambiae* |
| ERS1119031 | 2572STDY6370041 | AN0479-C | Cameroon | *An. gambiae* |
| ERS1119032 | 2572STDY6370042 | AN0480-C | Cameroon | *An. gambiae* |
| ERS1119033 | 2572STDY6370043 | AN0485-C | Cameroon | *An. gambiae* |
| ERS1119034 | 2572STDY6370044 | AN0486-C | Cameroon | *An. gambiae* |
| ERS1119035 | 2572STDY6370045 | AN0487-C | Cameroon | *An. gambiae* |
| ERS1119036 | 2572STDY6370046 | AN0488-C | Cameroon | *An. gambiae* |
| ERS1119037 | 2572STDY6370047 | AN0491-C | Cameroon | *An. gambiae* |
| ERS1119038 | 2572STDY6370048 | AN0492-C | Cameroon | *An. gambiae* |
| ERS1119039 | 2572STDY6370049 | AN0493-C | Cameroon | *An. coluzzii* |
| ERS1119040 | 2572STDY6370050 | AN0494-C | Cameroon | *An. coluzzii* |
| ERS1118969 | 2572STDY6370051 | AB0298-C | Burkina Faso | *An. gambiae* |
| ERS1118970 | 2572STDY6370052 | AB0299-C | Burkina Faso | *An. gambiae* |
| ERS1118971 | 2572STDY6370053 | AB0301-C | Burkina Faso | *An. gambiae* |
| ERS1118972 | 2572STDY6370054 | AB0302-C | Burkina Faso | *An. gambiae* |
| ERS1118973 | 2572STDY6370055 | AB0303-C | Burkina Faso | *An. gambiae* |
| ERS1118974 | 2572STDY6370056 | AB0309-C | Burkina Faso | *An. gambiae* |
| ERS1118975 | 2572STDY6370057 | AB0312-C | Burkina Faso | *An. gambiae* |
| ERS1118976 | 2572STDY6370058 | AB0313-C | Burkina Faso | *An. gambiae* |
| ERS1118977 | 2572STDY6370059 | AB0314-C | Burkina Faso | *An. gambiae* |
| ERS1118978 | 2572STDY6370060 | AB0315-C | Burkina Faso | *An. gambiae* |
| ERS1118979 | 2572STDY6370061 | AB0316-C | Burkina Faso | *An. gambiae* |
| ERS1118980 | 2572STDY6370062 | AB0317-C | Burkina Faso | *An. gambiae* |
| ERS1118981 | 2572STDY6370063 | AB0318-C | Burkina Faso | *An. coluzzii* |
| ERS1118982 | 2572STDY6370064 | AB0319-C | Burkina Faso | *An. coluzzii* |
| ERS1118983 | 2572STDY6370065 | AB0320-C | Burkina Faso | *An. coluzzii* |
| ERS1118984 | 2572STDY6370066 | AB0321-C | Burkina Faso | *An. coluzzii* |
| ERS1118985 | 2572STDY6370067 | AB0322-C | Burkina Faso | *unknown* |
| ERS1118986 | 2572STDY6370068 | AB0323-C | Burkina Faso | *An. coluzzii* |
| ERS1118987 | 2572STDY6370069 | AB0324-C | Burkina Faso | *An. coluzzii* |
| ERS1118988 | 2572STDY6370070 | AB0325-C | Burkina Faso | *An. coluzzii* |
| ERS1118967 | 2572STDY6369999 | AB0295-C | Cameroon | *An. gambiae* |
| ERS1118968 | 2572STDY6370000 | AB0297-C | Cameroon | *An. gambiae* |
| ERS1118991 | 2572STDY6370001 | AN0421-C | Cameroon | *An. gambiae* |
| ERS1118992 | 2572STDY6370002 | AN0423-C | Cameroon | *An. gambiae* |
| ERS1118993 | 2572STDY6370003 | AN0424-C | Cameroon | *An. gambiae* |

| | | | | |
|---|---|---|---|---|
| ERS1118994 | 2572STDY6370004 | AN0425-C | Cameroon | *An. gambiae* |
| ERS1118995 | 2572STDY6370005 | AN0426-C | Cameroon | *An. gambiae* |
| ERS1118996 | 2572STDY6370006 | AN0427-C | Cameroon | *An. gambiae* |
| ERS1118997 | 2572STDY6370007 | AN0428-C | Cameroon | *An. gambiae* |
| ERS1118998 | 2572STDY6370008 | AN0430-C | Cameroon | *An. gambiae* |
| ERS1118999 | 2572STDY6370009 | AN0431-C | Cameroon | *An. gambiae* |
| ERS1119000 | 2572STDY6370010 | AN0433-C | Cameroon | *An. coluzzii* |
| ERS1119001 | 2572STDY6370011 | AN0434-C | Cameroon | *An. gambiae* |
| ERS1119002 | 2572STDY6370012 | AN0435-C | Cameroon | *An. coluzzii* |
| ERS1119003 | 2572STDY6370013 | AN0436-C | Cameroon | *An. coluzzii* |
| ERS1119004 | 2572STDY6370014 | AN0437-C | Cameroon | *An. coluzzii* |
| ERS1119005 | 2572STDY6370015 | AN0438-C | Cameroon | *An. coluzzii* |
| ERS1119006 | 2572STDY6370016 | AN0440-C | Cameroon | *An. coluzzii* |
| ERS1119007 | 2572STDY6370017 | AN0442-C | Cameroon | *An. coluzzii* |
| ERS1119008 | 2572STDY6370018 | AN0443-C | Cameroon | *An. gambiae* |
| ERS1118965 | 2572STDY6369997 | AB0293-C | Cameroon | *An. gambiae* |
| ERS1118966 | 2572STDY6369998 | AB0294-C | Cameroon | *An. gambiae* |
| ERS1119011 | 2572STDY6370021 | AN0448-C | Cameroon | *An. gambiae* |
| ERS1119012 | 2572STDY6370022 | AN0449-C | Cameroon | *An. gambiae* |
| ERS1119013 | 2572STDY6370023 | AN0451-C | Cameroon | *An. gambiae* |
| ERS1119014 | 2572STDY6370024 | AN0453-C | Cameroon | *An. gambiae* |
| ERS1119015 | 2572STDY6370025 | AN0454-C | Cameroon | *An. gambiae* |
| ERS1119016 | 2572STDY6370026 | AN0455-C | Cameroon | *An. gambiae* |
| ERS1119017 | 2572STDY6370027 | AN0457-C | Cameroon | *An. gambiae* |
| ERS1119018 | 2572STDY6370028 | AN0458-C | Cameroon | *An. gambiae* |
| ERS1119019 | 2572STDY6370029 | AN0460-C | Cameroon | *An. gambiae* |
| ERS1119020 | 2572STDY6370030 | AN0461-C | Cameroon | *An. gambiae* |
| ERS1119021 | 2572STDY6370031 | AN0466-C | Cameroon | *An. gambiae* |
| ERS1119022 | 2572STDY6370032 | AN0468-C | Cameroon | *An. gambiae* |
| ERS1119023 | 2572STDY6370033 | AN0469-C | Cameroon | *An. gambiae* |
| ERS1119024 | 2572STDY6370034 | AN0470-C | Cameroon | *An. gambiae* |
| ERS1119025 | 2572STDY6370035 | AN0471-C | Cameroon | *An. gambiae* |
| ERS1119026 | 2572STDY6370036 | AN0474-C | Cameroon | *An. gambiae* |
| ERS1119027 | 2572STDY6370037 | AN0475-C | Cameroon | *An. gambiae* |
| ERS1119028 | 2572STDY6370038 | AN0476-C | Cameroon | *An. gambiae* |
| ERS1119042 | 2572STDY6370072 | AZ0253-C | Mali | *An. gambiae* |
| ERS1119043 | 2572STDY6370073 | AZ0254-C | Mali | *An. gambiae* |
| ERS1119044 | 2572STDY6370074 | AZ0255-C | Mali | *An. gambiae* |
| ERS1119045 | 2572STDY6370075 | AZ0256-C | Mali | *An. gambiae* |
| ERS1119046 | 2572STDY6370076 | AZ0257-C | Mali | *An. gambiae* |
| ERS1119047 | 2572STDY6370077 | AZ0258-C | Mali | *An. gambiae* |
| ERS1119048 | 2572STDY6370078 | AZ0259-C | Mali | *An. gambiae* |
| ERS1119049 | 2572STDY6370079 | AZ0260-C | Mali | *An. gambiae* |
| ERS1119050 | 2572STDY6370080 | AZ0261-C | Mali | *An. gambiae* |
| ERS1119051 | 2572STDY6370081 | AZ0262-C | Mali | *An. gambiae* |

| | | | | |
|---|---|---|---|---|
| ERS1119052 | 2572STDY6370082 | AZ0263-C | Mali | *An. gambiae* |
| ERS1119053 | 2572STDY6370083 | AZ0264-C | Mali | *An. coluzzii* |
| ERS1119056 | 2572STDY6370086 | AZ0267-C | Mali | *An. gambiae* |
| ERS1119057 | 2572STDY6370087 | AZ0268-C | Mali | *An. coluzzii* |
| ERS1119058 | 2572STDY6370088 | AZ0269-C | Mali | *An. coluzzii* |
| ERS1119059 | 2572STDY6370091 | AZ0270-C | Mali | *An. coluzzii* |
| ERS1119060 | 2572STDY6370092 | AZ0271-C | Mali | *An. coluzzii* |
| ERS1119061 | 2572STDY6370093 | AZ0272-C | Mali | *An. coluzzii* |
| ERS1119062 | 2572STDY6370094 | AZ0273-C | Mali | *An. coluzzii* |
| ERS1119063 | 2572STDY6370095 | AZ0274-C | Mali | *An. coluzzii* |
| ERS1119064 | 2572STDY6370096 | AZ0275-C | Mali | *An. coluzzii* |
| ERS1119065 | 2572STDY6370097 | AZ0276-C | Mali | *An. coluzzii* |
| ERS1119066 | 2572STDY6370098 | AZ0277-C | Mali | *An. coluzzii* |
| ERS1119067 | 2572STDY6370099 | AZ0278-C | Mali | *An. coluzzii* |
| ERS1119068 | 2572STDY6370100 | AZ0279-C | Mali | *An. gambiae* |
| ERS1119069 | 2572STDY6370101 | AZ0280-C | Mali | *An. coluzzii* |
| ERS1119070 | 2572STDY6370102 | AZ0281-C | Mali | *An. coluzzii* |
| ERS1119071 | 2572STDY6370103 | AZ0282-C | Mali | *An. coluzzii* |
| ERS1119072 | 2572STDY6370104 | AZ0283-C | Mali | *An. gambiae* |
| ERS1502572 | 4431STDY6672871 | VBS02003 | Cameroon | *An. gambiae* |
| ERS1502573 | 4431STDY6672872 | VBS02004 | Cameroon | *An. gambiae* |
| ERS1502574 | 4431STDY6672873 | VBS02005 | Cameroon | *An. gambiae* |
| ERS1502575 | 4431STDY6672874 | VBS02006 | Cameroon | *An. gambiae* |
| ERS1502576 | 4431STDY6672875 | VBS02007 | Cameroon | *An. gambiae* |
| ERS1502577 | 4431STDY6672876 | VBS02008 | Cameroon | *An. gambiae* |
| ERS1502578 | 4431STDY6672877 | VBS02009 | Cameroon | *An. gambiae* |
| ERS1502579 | 4431STDY6672878 | VBS02010 | Cameroon | *An. gambiae* |
| ERS1502580 | 4431STDY6672879 | VBS02011 | Cameroon | *An. gambiae* |
| ERS1502581 | 4431STDY6672880 | VBS02012 | Cameroon | *An. gambiae* |
| ERS1502582 | 4431STDY6672881 | VBS02013 | Cameroon | *An. gambiae* |
| ERS1502583 | 4431STDY6672882 | VBS02014 | Cameroon | *An. gambiae* |
| ERS1502584 | 4431STDY6672883 | VBS02015 | Cameroon | *An. gambiae* |
| ERS1502585 | 4431STDY6672884 | VBS02016 | Cameroon | *An. gambiae* |
| ERS1502586 | 4431STDY6672885 | VBS02017 | Cameroon | *An. gambiae* |
| ERS1502587 | 4431STDY6672886 | VBS02018 | Cameroon | *An. gambiae* |
| ERS1502589 | 4431STDY6672888 | VBS02020 | Cameroon | *An. gambiae* |
| ERS1502590 | 4431STDY6672889 | VBS02021 | Cameroon | *An. gambiae* |
| ERS1502591 | 4431STDY6672890 | VBS02022 | Cameroon | *An. gambiae* |
| ERS1502592 | 4431STDY6672891 | VBS02023 | Cameroon | *An. gambiae* |
| ERS1502593 | 4431STDY6672892 | VBS02024 | Cameroon | *An. gambiae* |
| ERS1502594 | 4431STDY6672893 | VBS02025 | Cameroon | *An. gambiae* |
| ERS1502595 | 4431STDY6672894 | VBS02026 | Cameroon | *An. gambiae* |
| ERS1502596 | 4431STDY6672895 | VBS02027 | Cameroon | *An. gambiae* |
| ERS1502597 | 4431STDY6672896 | VBS02028 | Cameroon | *An. gambiae* |
| ERS1502598 | 4431STDY6672897 | VBS02029 | Cameroon | *An. gambiae* |

| | | | | |
|---|---|---|---|---|
| ERS1502599 | 4431STDY6672898 | VBS02030 | Cameroon | *An. gambiae* |
| ERS1502600 | 4431STDY6672899 | VBS02031 | Cameroon | *An. gambiae* |
| ERS1502601 | 4431STDY6672900 | VBS02032 | Cameroon | *An. gambiae* |
| ERS1502602 | 4431STDY6672901 | VBS02033 | Cameroon | *An. gambiae* |
| ERS1502603 | 4431STDY6672902 | VBS02034 | Cameroon | *An. gambiae* |
| ERS1502604 | 4431STDY6672903 | VBS02035 | Cameroon | *An. gambiae* |
| ERS1502605 | 4431STDY6672904 | VBS02036 | Cameroon | *An. gambiae* |
| ERS1502606 | 4431STDY6672905 | VBS02037 | Cameroon | *An. gambiae* |
| ERS1502607 | 4431STDY6672906 | VBS02038 | Cameroon | *An. gambiae* |
| ERS1502608 | 4431STDY6672907 | VBS02039 | Cameroon | *An. gambiae* |
| ERS1502609 | 4431STDY6672908 | VBS02040 | Cameroon | *An. gambiae* |
| ERS1502610 | 4431STDY6672909 | VBS02041 | Cameroon | *An. gambiae* |
| ERS1502611 | 4431STDY6672910 | VBS02042 | Cameroon | *An. gambiae* |
| ERS1502612 | 4431STDY6672911 | VBS02043 | Cameroon | *An. gambiae* |
| ERS1502613 | 4431STDY6672912 | VBS02044 | Cameroon | *An. gambiae* |
| ERS1502614 | 4431STDY6672913 | VBS02045 | Cameroon | *An. gambiae* |
| ERS1502615 | 4431STDY6672914 | VBS02046 | Cameroon | *An. gambiae* |
| ERS1502616 | 4431STDY6672915 | VBS02047 | Cameroon | *An. gambiae* |
| ERS1502617 | 4431STDY6672916 | VBS02048 | Cameroon | *An. gambiae* |
| ERS1502618 | 4431STDY6672917 | VBS02049 | Mali | *An. coluzzii* |
| ERS1502619 | 4431STDY6672918 | VBS02050 | Mali | *An. coluzzii* |
| ERS1502620 | 4431STDY6672919 | VBS02051 | Mali | *An. coluzzii* |
| ERS1502621 | 4431STDY6672920 | VBS02052 | Mali | *An. coluzzii* |
| ERS1502622 | 4431STDY6672921 | VBS02053 | Mali | *An. coluzzii* |
| ERS1502623 | 4431STDY6672922 | VBS02054 | Mali | *An. coluzzii* |
| ERS1502624 | 4431STDY6672923 | VBS02055 | Mali | *An. coluzzii* |
| ERS1502626 | 4431STDY6672925 | VBS02057 | Mali | *An. coluzzii* |
| ERS1502627 | 4431STDY6672926 | VBS02058 | Mali | *An. coluzzii* |
| ERS1502628 | 4431STDY6672927 | VBS02059 | Mali | *An. coluzzii* |
| ERS1502629 | 4431STDY6672928 | VBS02060 | Mali | *An. coluzzii* |
| ERS1502631 | 4431STDY6672930 | VBS02062 | Mali | *An. coluzzii* |
| ERS1502632 | 4431STDY6672931 | VBS02063 | Mali | *An. coluzzii* |
| ERS1502633 | 4431STDY6672932 | VBS02064 | Mali | *An. coluzzii* |
| ERS1502634 | 4431STDY6672933 | VBS02065 | Mali | *An. coluzzii* |
| ERS1502635 | 4431STDY6672934 | VBS02066 | Mali | *An. coluzzii* |
| ERS1502636 | 4431STDY6672935 | VBS02067 | Mali | *An. coluzzii* |
| ERS1502637 | 4431STDY6672936 | VBS02068 | Mali | *An. coluzzii* |
| ERS1502638 | 4431STDY6672937 | VBS02069 | Mali | *An. coluzzii* |
| ERS1502639 | 4431STDY6672938 | VBS02070 | Mali | *An. coluzzii* |
| ERS1502640 | 4431STDY6672939 | VBS02071 | Mali | *An. coluzzii* |
| ERS1502641 | 4431STDY6672940 | VBS02072 | Mali | *An. coluzzii* |
| ERS1502642 | 4431STDY6672941 | VBS02073 | Mali | *An. coluzzii* |
| ERS1625214 | 4431STDY6772804 | VBS02076 | Mali | *An. gambiae* |
| ERS1625215 | 4431STDY6772805 | VBS02077 | Mali | *An. gambiae* |
| ERS1625216 | 4431STDY6772806 | VBS02083 | Mali | *An. gambiae* |

| | | | | |
|---|---|---|---|---|
| ERS1625217 | 4431STDY6772807 | VBS02085 | Mali | *An. gambiae* |
| ERS1625218 | 4431STDY6772808 | VBS02088 | Mali | *An. gambiae* |
| ERS1625219 | 4431STDY6772809 | VBS02090 | Mali | *An. gambiae* |
| ERS1625220 | 4431STDY6772810 | VBS02092 | Mali | *An. gambiae* |
| ERS1625221 | 4431STDY6772811 | VBS02093 | Mali | *An. gambiae* |
| ERS1625222 | 4431STDY6772812 | VBS02094 | Mali | *An. gambiae* |
| ERS1625223 | 4431STDY6772813 | VBS02095 | Mali | *An. gambiae* |
| ERS1625224 | 4431STDY6772814 | VBS02096 | Mali | *An. gambiae* |
| ERS1625225 | 4431STDY6772815 | VBS02097 | Mali | *An. gambiae* |
| ERS1625227 | 4431STDY6772817 | VBS02100 | Mali | *An. gambiae* |
| ERS1625228 | 4431STDY6772818 | VBS02101 | Mali | *An. gambiae* |
| ERS1625229 | 4431STDY6772819 | VBS02103 | Mali | *An. gambiae* |
| ERS1625230 | 4431STDY6772820 | VBS02104 | Mali | *An. gambiae* |
| ERS1625231 | 4431STDY6772821 | VBS02106 | Mali | *An. gambiae* |
| ERS1625232 | 4431STDY6772822 | VBS02107 | Mali | *An. gambiae* |
| ERS1625233 | 4431STDY6772823 | VBS02109 | Mali | *An. gambiae* |
| ERS1625234 | 4431STDY6772824 | VBS02110 | Mali | *An. gambiae* |
| ERS1625236 | 4431STDY6772826 | VBS02112 | Mali | *An. gambiae* |
| ERS1625237 | 4431STDY6772827 | VBS02113 | Mali | *An. gambiae* |
| ERS1625238 | 4431STDY6772828 | VBS02114 | Mali | *An. gambiae* |
| ERS1625239 | 4431STDY6772829 | VBS02115 | Mali | *An. gambiae* |
| ERS1625241 | 4431STDY6772831 | VBS02119 | Mali | *An. gambiae* |
| ERS1625242 | 4431STDY6772832 | VBS02121 | Mali | *An. gambiae* |
| ERS1625243 | 4431STDY6772833 | VBS02127 | Mali | *An. gambiae* |
| ERS1625244 | 4431STDY6772834 | VBS02128 | Mali | *An. gambiae* |
| ERS1625245 | 4431STDY6772835 | VBS02129 | Mali | *An. gambiae* |
| ERS1625246 | 4431STDY6772836 | VBS02130 | Mali | *An. gambiae* |
| ERS1625247 | 4431STDY6772837 | VBS02132 | Mali | *An. gambiae* |
| ERS1625248 | 4431STDY6772838 | VBS02133 | Mali | *An. gambiae* |

**Table S2**. Ag1000G and Vector Observatory *An. gambiae* and *An. coluzzii* mosquitoes used for computational karyotyping.

| | Number of Specimens (inversions karyotyped) | | |
|---|---|---|---|
| **Source Country** | ***An. coluzzii*** | ***An. gambiae*** | **(Inferred) Hybrids** |
| Angola[1, 2] | 78 (*a, b, c, u*) | 0 | 0 |
| Burkina Faso[1, 2, 3] | 82 (*a, b, c, u*) | 104 (*a, b, c, d, j, u*) | 1 (*a, b, u*) |
| Cameroon[1, 2, 3] | 9 (*a, b, c, u*) | 389 (*a, b, c, d, j, u*) | 0 |
| Cote d'Ivoire[2] | 71 (*a, b, c, u*) | 0 | 0 |
| Equatorial Guinea[2] | 0 | 9 (*a, b, c, d, j, u*) | 0 |
| France (Mayotte)[2] | 0 | 24 (*a, b, c, d, u*) | 0 |
| Gabon[1, 2] | 0 | 69 (*a, b, c, d, j, u*) | 0 |
| The Gambia[2] | 0 | 0 | 65 (*a, b*) |
| Ghana[2] | 55 (*a, b, c, u*) | 12 (*a, b, c, d, j, u*) | 0 |
| Guinea[1,2] | 4 (*a, b, c, u*) | 39 (*a, b, c, d, j, u*) | 1 (*a, b, u*) |
| Guinea-Bissau[1, 2] | 4 (*a, b*) | 66 (*a, b*) | 21 (*a, b*) |
| Kenya[1, 2] | 0 | 48 (*a*) | 0 |
| Mali[3] | 38 (*a, b, c, u*) | 46 (*a, b, c, d, j, u*) | 0 |
| Uganda[1, 2] | 0 | 112 (*a, b, c, d, j, u*) | 0 |
| **Total** | **341** | **918** | **88** |

[1]Ag1000G phase 1 AR3 data release (https://www.malariagen.net/data/ag1000g-phase1-ar3)
[2]Ag1000G phase 2 AR1 data release (https://www.malariagen.net/data/ag1000g-phase-2-ar1)
[3]Ag1000G phase 3 and Vector Observatory data, available from (https://figshare.com/projects/Data_for_In_silico_karyotyping_of_chromosomally_polymorphic_malaria_mosquitoes_in_the_Anopheles_gambiae_complex_/65522) under the Ag1000G terms of use: https://www.malariagen.net/data/terms-use/ag1000g-terms-use

**Table S3**. *An. gambiae* AgamP4 reference coordinates taken as inversion boundaries for the purpose of PCA and computational karyotyping.

| **Inversion** | **Inversion boundaries for karyotyping** |
|---|---|
| 2L*a* | 20524058-42165532[1] |
| 2R*j* | 3262186-15750717[2] |
| 2R*b* | 19023925-26758676[3] |
| 2R*c* | 26750000-31473100[4] |
| 2R*d* | 31495381-42375004[5] (Computational karyotyping)<br>41000000-42375004 (PCA karyotyping) |
| 2R*u* | 31473000-35505236[4] |

[1]Sharakhov, I.V., B.J. White, M.V. Sharakhova, J. Kayondo, N.F. Lobo *et al.*, 2006 Breakpoint structure reveals the unique origin of an interspecific chromosomal inversion (2La) in the *Anopheles gambiae* complex. *Proc Natl Acad Sci U S A* 103:6258-6262.
[2]Coulibaly, M.B., N.F. Lobo, M.C. Fitzpatrick, M. Kern, O. Grushko *et al.*, 2007 Segmental duplication implicated in the genesis of inversion 2R*j* of *Anopheles gambiae*. *PloS ONE* 2:e849.
[3]Lobo, N.F., D.M. Sangare, A.A. Regier, K.R. Reidenbach, D.A. Bretz *et al.*, 2010 Breakpoint structure of the *Anopheles gambiae* 2R*b* chromosomal inversion. *Malar J* 9:293.
[4]Sangare, D.M., 2007 Breakpoint analysis of the *Anopheles gambiae s.s.* chromosome 2R*b*, 2R*c*, and 2R*u* inversions in *PhD Thesis, Graduate Program in Biological Sciences, University of Notre Dame*. University of Notre Dame, Notre Dame, IN.
[5]Corbett-Detig, R., I. Said, M. Calzetta, M. Genetti, J. McBroome *et al.*, 2019 Fine-mapping complex inversion breakpoints and investigating somatic pairing in the *Anopheles gambiae* species complex using proximity-ligation sequencing. *BioRxiv doi:* *https://doi.org/10.1101/662114*

**Table S4.** Number of Ag1000G *An. gambiae* and *An. coluzzii* mosquitoes with PCA karyotype assignments.

| | **Karyotype[1] based on PCA** | | | **Total karyotyped** | **Not karyotyped by PCA** |
|---|---|---|---|---|---|
| | **0** | **1** | **2** | | |
| 2L*a* | 502 | 353 | 492 | 1347 | 0 |
| 2R*j* | 731 | 4 | 45 | 780 | 567 (*An. coluzzii*, The Gambia, Guinea Bissau, Kenya, Mayotte) |
| 2R*b* | 727 | 331 | 241 | 1299 | 48 (Kenya) |
| 2R*c* | 962 | 89 | 92 | 1143 | 204 (The Gambia, Guinea Bissau, Kenya) |
| 2R*u* | 1040 | 47 | 56 | 1143 | 204 (The Gambia, Guinea Bissau, Kenya) |
| 2R*d* | 750 | 34 | 20 | 804 | 543 (*An. coluzzii*, The Gambia, Guinea Bissau, Kenya) |

[1]Adapted from Touré et al (1998), '0' represents the homokaryotypic standard, '1' the heterokaryotype, and '2' the homokaryotypic inverted genotype.

**Table S5.** Performance of computational karyotyping on low coverage re-sequencing data for inversions 2L*a*, 2R*b*, 2R*c*, and 2R*u*.

| | | **2L*a*** | | | | **2R*b*** | | | | **2Rc** | | | | **2R*u*** | | | |
|---|---|---|---|---|---|---|---|---|---|---|---|---|---|---|---|---|---|
| **Specimen ID** | **Cov.** | **CYT** | **TAG** | **# tags** | **tags matching comp. score (%)** | **CYT** | **TAG** | **# tags** | **tags matching comp. score (%)** | **CYT** | **TAG 2Rc-gam** | **# tags** | **tags matching comp. score (%)** | **CYT** | **TAG** | **# tags** | **tags matching comp. score (%)** |
| **BAMAKO**[1] | | | | | | | | | | | | | | | | | |
| KL0218 | 9.3 | 2 | 2.0 | 10 | 100.0 | 2 | 1.6 | 42 | 66.7 | 2 | 1.0 | 2 | 100.0 | 2 | 1.8 | 17 | 82.4 |
| KL0220 | 9.4 | 2 | 1.8 | 12 | 83.3 | 0 | 0.2 | 42 | 85.7 | 2 | 1.3 | 3 | 66.7 | 2 | 1.6 | 15 | 60.0 |
| KL0231 | 9.8 | 2 | 1.8 | 11 | 81.8 | 0 | 0.2 | 44 | 86.4 | 2 | 1.0 | 3 | 33.3 | 2 | 1.6 | 17 | 58.8 |
| KL0333 | 9.0 | N/A | 1.9 | 11 | 90.9 | 0 | 0.2 | 42 | 83.3 | 2 | 0.7 | 3 | 33.3 | 2 | 1.6 | 16 | 62.5 |
| KL0341 | 9.8 | N/A | 1.9 | 11 | 90.9 | 0 | 0.2 | 44 | 84.1 | 2 | 1.0 | 3 | 33.3 | 2 | 1.7 | 18 | 66.7 |
| KL0370 | 10.3 | N/A | 1.9 | 12 | 91.7 | 2 | 1.7 | 45 | 71.1 | 2 | 1.0 | 3 | 33.3 | 2 | 1.7 | 18 | 72.2 |
| KL0671 | 8.9 | N/A | 1.8 | 11 | 81.8 | 2 | 1.6 | 38 | 63.2 | 2 | 0.3 | 3 | 66.7 | 2 | 1.6 | 17 | 76.5 |
| KL0899 | 10.1 | N/A | 1.8 | 11 | 81.8 | 0 | 0.3 | 38 | 73.7 | 2 | 0.7 | 3 | 33.3 | 2 | 1.7 | 16 | 68.8 |
| ***An. coluzzii***[2] | | | | | | | | | | | **2Rc-col** | | | | | | |
| 02SEL85 | 20 | 2 | 2.0 | 197 | 100.0 | 2 | 1.0 | 330 | 81.8 | 2 | 1.0 | 55 | 92.7 | 0 | 0.0 | 164 | 98.2 |
| 04SEL18 | 13 | 2 | 2.0 | 196 | 99.0 | 2 | 1.8 | 322 | 79.5 | 2 | 1.8 | 53 | 84.9 | 0 | 0.0 | 143 | 97.2 |
| O10SEL160 | 14 | 1 | 2.0 | 197 | 99.5 | 2 | 1.8 | 330 | 83.9 | 0 | 1.8 | 49 | 83.7 | 0 | 0.1 | 148 | 94.6 |
| 04SEL021 | 4 | 1 | 1.1 | 108 | 30.6 | 1 | 0.9 | 160 | 24.4 | 1 | 0.9 | 28 | 35.7 | 0 | 0.1 | 52 | 92.3 |
| 2012SEL002 | 10 | 2 | 2.0 | 157 | 100.0 | 2 | 1.8 | 233 | 85.0 | 2 | 1.7 | 29 | 75.9 | 0 | 0.1 | 58 | 93.1 |
| 2012SEL003 | 14 | 2 | 2.0 | 167 | 99.4 | 0 | 0.2 | 233 | 85.8 | 0 | 0.1 | 35 | 91.4 | 0 | 0.1 | 83 | 95.2 |
| 2012SEL006 | 10 | 2 | 2.0 | 159 | 100.0 | 0 | 0.1 | 226 | 89.8 | 0 | 0.0 | 30 | 96.7 | 2 | 1.1 | 120 | 65.0 |
| 2012SEL009 | 18 | 2 | 2.0 | 188 | 100.0 | 0 | 0.2 | 304 | 84.9 | 0 | 0.2 | 50 | 86.0 | 2 | 1.0 | 156 | 87.2 |
| 2012SEL013 | 23 | 2 | 2.0 | 192 | 100.0 | 1 | 1.0 | 315 | 77.8 | 1 | 1.0 | 51 | 80.4 | 1 | 1.0 | 154 | 81.2 |
| 010sel134 | 17 | 1 | 2.0 | 194 | 100.0 | 2 | 0.2 | 326 | 85.9 | 0 | 0.1 | 55 | 92.7 | 0 | 1.0 | 166 | 92.2 |
| 2012sel012 | 8 | 2 | 2.0 | 188 | 98.9 | 0 | 0.1 | 281 | 87.2 | 0 | 0.1 | 46 | 93.5 | 0 | 0.1 | 99 | 92.9 |
| 2012sel029 | 9 | 2 | 2.0 | 190 | 99.5 | 2 | 1.8 | 283 | 82.7 | 2 | 1.9 | 41 | 92.7 | 0 | 0.0 | 98 | 98.0 |
| 2012sel063 | 15 | 2 | 2.0 | 193 | 99.5 | 2 | 1.8 | 329 | 81.5 | 2 | 1.7 | 53 | 75.5 | 0 | 0.0 | 144 | 95.8 |
| 04SEL02 | 5 | 2 | 2.0 | 154 | 100.0 | 0 | 1.7 | 208 | 81.3 | 0 | 0.9 | 32 | 53.1 | 1 | 0.1 | 57 | 94.7 |
| 04SEL14 | 40 | 2 | 2.0 | 196 | 100.0 | 2 | 1.8 | 332 | 88.0 | 2 | 1.8 | 56 | 76.8 | 0 | 0.1 | 168 | 94.6 |
| 04SEL84 | 26 | 2 | 2.0 | 196 | 100.0 | 1 | 1.0 | 332 | 81.9 | 1 | 0.9 | 56 | 89.3 | 0 | 0.0 | 166 | 95.8 |
| 04SEL91 | 66 | 2 | 2.0 | 198 | 99.0 | 1 | 1.0 | 334 | 82.9 | 1 | 1.0 | 56 | 92.9 | 0 | 0.0 | 169 | 98.2 |

Cov, mean sequencing coverage; CYT, cytogenetic karyotype; TAG, computational karyotype score; # tags, number of tags scored. Gray cells highlight mismatches between cytogenetic and computational karyotype.

[1]Love, R. R., A. M. Steele, M. B. Coulibaly, S. F. Traore, S. J. Emrich *et al.*, 2016 Chromosomal inversions and ecotypic differentiation in *Anopheles gambiae*: the perspective from whole-genome sequencing. *Mol. Ecol.* 25: 5889-5906.

[2]Main, B. J., Y. Lee, T. C. Collier, L. C. Norris, K. Brisco *et al.*, 2015 Complex genome evolution in *Anopheles coluzzi*i associated with increased insecticide usage in Mali. *Mol. Ecol.* 24: 5145-5157

**Table S6.** Performance of computational karyotyping on low coverage re-sequencing data for inversions 2R*d* and 2R*j*

| | | **2R*d*** | | | | **2R*j*** | | | |
|---|---|---|---|---|---|---|---|---|---|
| **Specimen ID** | **Cov.** | **CYT** | **TAG 2R*d*-gam** | **# tags** | **tags matching comp. score (%)** | **CYT** | **TAG 2R*j*-gam** | **# tags** | **tags matching comp. score (%)** |
| **BAMAKO**[1] | | | | | | | | | |
| KL0218 | 9.3 | 0 | 0.3 | 24 | 79.2 | 2 | 1.5 | 11 | 63.6 |
| KL0220 | 9.4 | 0 | 0.1 | 25 | 92.0 | 2 | 1.8 | 11 | 81.8 |
| KL0231 | 9.8 | 0 | 0.2 | 26 | 84.6 | 2 | 1.8 | 13 | 76.9 |
| KL0333 | 9.0 | 0 | 0.3 | 23 | 78.3 | 2 | 1.8 | 13 | 76.9 |
| KL0341 | 9.8 | 0 | 0.3 | 24 | 79.2 | 2 | 1.8 | 12 | 75.0 |
| KL0370 | 10.3 | 0 | 0.1 | 25 | 92.0 | 2 | 1.8 | 12 | 83.3 |
| KL0671 | 8.9 | 0 | 0.3 | 23 | 78.3 | 2 | 1.7 | 7 | 71.4 |
| KL0899 | 10.1 | 0 | 0.3 | 25 | 76.0 | 2 | 1.8 | 12 | 75.0 |

Cov, mean sequencing coverage; CYT, cytogenetic karyotype; TAG, computational karyotype score; # tags, number of tags scored. Gray cells highlight mismatches between cytogenetic and computational karyotype.

[1]Love, R. R., A. M. Steele, M. B. Coulibaly, S. F. Traore, S. J. Emrich *et al.*, 2016 Chromosomal inversions and ecotypic differentiation in *Anopheles gambiae*: the perspective from whole-genome sequencing. *Mol. Ecol.* 25: 5889-5906.

**Table S7.** Performance of tag SNPs relative to cytogenetic karyotype assignments when used to genotype heterospecific or admixed mosquito samples.

| | | | | **Specimens with discrepancies** | | |
|---|---|---|---|---|---|---|
| **Inversion Tags** | **Heterospecific or admixed sample** | **CYT** | **Specimens (N)** | **Mismatch CYT-TAG (%)** | **Match TAG-PCA (%)** | **No. tag SNPs scored (% matching TAG)** |
| **2R*j*-*gambiae*** | *coluzzii* | 0 | 54 | 0 (0) | – | – |
| | G-B[1] | 0 | 33 | 0 (0) | – | – |
| **2R*c*-gambiae[2]** | BAMAKO | 2 | 45 | 27 (60.0) | 0 (0) | 36-49 (14.3-71.4) |
| **2R*c*-gam+col[3]** | G-B[1] | 0 | 33 | 0 (0) | – | – |
| **2R*d*-*gambiae*** | *coluzzii* | 0 | 48 | 0 (0) | N/A[4] | – |
| | | 1 | 6 | 6 (100) | N/A[4] | 139-147 (94.6-95.9) |
| | G-B[1] | 0 | 19 | 0 (0) | N/A[5] | – |
| | | 2 | 14 | 14 (100) | N/A[5] | 146-147 (76.7-96.6) |
| **2R*u*** | G-B[1] | 0 | 33 | 0 (0) | N/A[5] | – |

CYT, cytogenetic genotype; TAG, computational genotype; PCA, genotype inferred by PCA.
[1]Specimens from Guinea-Bissau (G-B), where elevated *An. gambiae-An. coluzzii* admixture occurs.
[2]2Rc tags designed to genotype *An. gambiae* excluding BAMAKO
[3]Results were the same using either 2Rc-gambiae tags or 2Rc-coluzzii tags when genotyping specimens from Guinea-Bissau.
[4]*An. coluzzii* could not be karyotyped by PCA for 2R*d*
[5]Specimens from Guinea-Bissau could not be karyotyped by PCA for 2R*d* or 2R*u*

**Table S8.** Software used.

| Name | Most recent version used | Reference or source |
|---|---|---|
| *Command line tools* | | |
| bcftools | 1.4-6-g5e49659 | http://www.htslib.org/doc/bcftools.html |
| tabix | 0.2.5 (r1005) | https://github.com/samtools/htslib |
| *Python packages* | | |
| anhima | 0.11.2 | https://github.com/alimanfoo/anhima, commit d9eed2d |
| cartopy | 0.17.1.dev168+ | Cartopy. Met Office. git@github.com:SciTools/cartopy.git. 2019-06-10. 1942c7b. |
| collections | *python 3.7.3* | |
| datetime | *python 3.7.3* | |
| h5py | 2.9.0 | https://www.h5py.org/ |
| ingenos | 0.1 | https://github.com/rrlove/ingenos |
| itertools | *python 3.7.3* | |
| matplotlib | 3.1.0 | J. D. Hunter, "Matplotlib: A 2D Graphics Environment", Computing in Science & Engineering, vol. 9, no. 3, pp. 90-95, 2007.<br><br>DOI:10.5281/zenodo.2893252 |
| mpl_toolkits | *accompanies matplotlib 3.1.0* | |
| numpy | 1.16.3 | Stéfan van der Walt, S. Chris Colbert and Gaël Varoquaux. **The NumPy Array: A Structure for Efficient Numerical Computation**, Computing in Science & Engineering, **13**, 22-30 (2011), DOI:10.1109/MCSE.2011.37 (publisher link) |
| os | *python 3.7.3* | |
| pandas | 0.24.2 | Wes McKinney. **Data Structures for Statistical Computing in Python**, Proceedings of the 9th Python in Science Conference, 51-56 (2010) (publisher link) |
| re | 2.2.1 | *accompanies python 3.7.3* |
| rpy2 | 2.9.5 | https://rpy2.readthedocs.io/en/version_2.8.x/ |
| scikit-allel | 1.2.0 | DOI:10.5281/zenodo.3238280 |
| scikit-learn | 0.21.1 | Fabian Pedregosa, Gaël Varoquaux, Alexandre Gramfort, Vincent Michel, Bertrand Thirion, Olivier Grisel, Mathieu Blondel, Peter Prettenhofer, Ron Weiss, Vincent Dubourg, Jake Vanderplas, Alexandre Passos, David Cournapeau, Matthieu Brucher, Matthieu Perrot, Édouard Duchesnay. **Scikit-learn: Machine Learning in Python**, Journal of Machine Learning Research, **12**, 2825-2830 (2011) (publisher link) |
| scipy | 1.3.0 | Jones E, Oliphant E, Peterson P, *et al.* **SciPy: Open Source Scientific Tools for Python**, 2001-, http://www.scipy.org/ [Online; accessed 2019-06-18]. |
| seaborn | 0.7.1 | DOI:10.5281/zenodo.54844 |
